# Structures of Fab-stabilized CHIP reveal a conformational switch important in E3 ligase and chaperone functions

**DOI:** 10.1101/2025.05.15.654159

**Authors:** Aparna Unnikrishnan, Dong hee Chung, Emily J. Connelly, Shih-Wei Chuo, Shweta Devi, Aye C. Thwin, Cory M. Nadel, Eric Tse, Jason E. Gestwicki, Charles S. Craik, Daniel R. Southworth

## Abstract

Carboxyl terminus of Hsc70-interacting protein (CHIP/STUB1) is a U-box E3 ligase essential for protein quality control, targeting misfolded or damaged proteins for clearance and conducting chaperone-like functions by suppressing aggregation of proteins, including tau. The previous structure of full-length CHIP identified an asymmetric homodimer in which one U-box is occluded from E2 binding, indicating an unusual half-of-sites activity. However, the flexibility of CHIP has complicated efforts to further characterize its structure and function. Here we leverage two CHIP-targeting fragment antigen-binding (Fab) antibodies to solve structures by cryo-EM. We identify one Fab binds to the CHIP U-box via interactions mimicking E2 contacts and stabilizes three distinct CHIP dimer states, revealing an asymmetric-to-symmetric conformational switch that would enable both U-box domains to be accessible for E2 binding. Conversely, the second Fab targets CHIP’s coiled-coil domains, stabilizing the asymmetric dimer with a single accessible U-box. Remarkably, the Fabs exhibit opposing effects on CHIP’s inhibition of tau aggregation, wherein binding to coiled-coil domains abolishes inhibition of aggregation, while binding to the U-box greatly potentiates this activity. Together, this work reveals how CHIP conformational states and binding interfaces may regulate ubiquitination cycles and chaperone-like functions.

## Introduction

The carboxyl terminus of Hsc70-interacting protein (CHIP/STUB1) is an E3 ligase that serves as a key co-chaperone regulator of protein homeostasis (proteostasis)^1,2^. CHIP has critical roles in pro-folding and pro-degradation pathways for many substrates, including proteotoxic forms of the microtubule associated protein tau, which is implicated in many “tauopathy” neurodegenerative diseases^1–6^. In tauopathies including Alzheimer’s Disease, CHIP co-localizes with tau deposits in the brain^5^. CHIP knockout mice show elevated levels of phosphorylated tau (p-tau) and reduced poly-ubiquitinated tau^7^, while CHIP overexpression accelerates degradation of p-tau^8,9^. Under non-stress conditions, CHIP activity regulates lifespan in worms and flies, where it promotes monoubiquitination of the insulin receptor for its endocytic-lysosomal turnover ^10,11^. Recently, CHIP has also been shown to have chaperone-like activities on its own, binding to p-tau and suppressing its aggregation^12^. Nonetheless, the structural states and mechanisms that enable CHIP’s diverse functions in ubiquitination and chaperone activity are not yet clear.

CHIP contains a C-terminal U-box domain that interacts with specific E2 ubiquitin conjugating enzymes, including Ubc13^13^, UbcH5^14^ and Ube2w^15^ to mediate ubiquitin transfer to substrates^16,17^. CHIP is an obligate homodimer^18^ with U-box domains interacting through well-conserved hydrophobic residues that pack to form a high-affinity parallel dimer interface, resulting in E2 binding sites that are positioned on either side of the CHIP dimer^13^. CHIP also contains an N-terminal tetratricopeptide repeat (TPR) domain that binds the C-terminal EEVD residues of cytosolic molecular chaperones: heat shock protein 70 (Hsp70)^19^ and 90 (Hsp90)^20^. This interaction is thought to be important for “triage” and degradation of misfolded chaperone-bound substrates^5,19–21^. The TPR domain also interacts with EEVD-like motifs in some proteins, circumventing the need for chaperones^12,22–24^. Finally, CHIP contains a central coiled-coil (CC) domain comprised of helices α7-8 (also referred to as “helical hairpin” or “HH”) that connects the TPR and U-box domains and makes additional dimerization contacts as a four-helix bundle^13^.

Studies of CHIP and its truncated forms have shown conflicting reports of either asymmetric^13^ or symmetric dimer conformations^14^ and the potential roles of these structural states are not yet clear. Full-length murine CHIP crystal structure with a bound EEVD peptide adopts an unexpected asymmetric conformation in which one protomer contains a “bent” (or “broken”) CC domain and a TPR domain that is positioned adjacent to the U-box in a manner that occludes E2 interactions (here referred to as “occluding TPR” or “TPR_o_”) (Fig. 1a, b)^13^. Conversely, the adjacent protomer contains an intact “straight” CC domain in which α7 is rigidly connected to TPR helix α6 resulting in a rotated position of the TPR domain (here referred to as “rotated TPR” or “TPR_r_”) that exposes the E2 binding site on the U-box^13^. Thus, this asymmetric CHIP dimer conformation has a single site that is accessible for E2 binding and Ub transfer. Indeed, binding studies with uncharged E2 indicated only a single E2 is capable of binding the CHIP dimer^13^. Thus, the CHIP dimer is proposed to adopt a partially auto-inhibited state with a sub-optimal “half-of-sites” activity for ubiquitination, highlighting questions about the functional role of this structure. Notably, when the TPR domains are fully truncated (e.g., CC-U-box), the protein still forms dimers, but they adopt a symmetrical CC (both straight) conformation, with U-box domains binding E2s with a 2:2 stoichiometry, displaying potential full-site reactivity^13,14^. This has also been reported by NMR studies using U-box dimers^25^. Moreover, molecular dynamic (MD) simulations have indicated asymmetric-to-symmetric dimer conversions may occur^26^. However, it is unclear how both E2 binding sites in the U-box dimer may become accessible or coordinate during CHIP-driven cycles of ubiquitination.

**Fig. 1.**
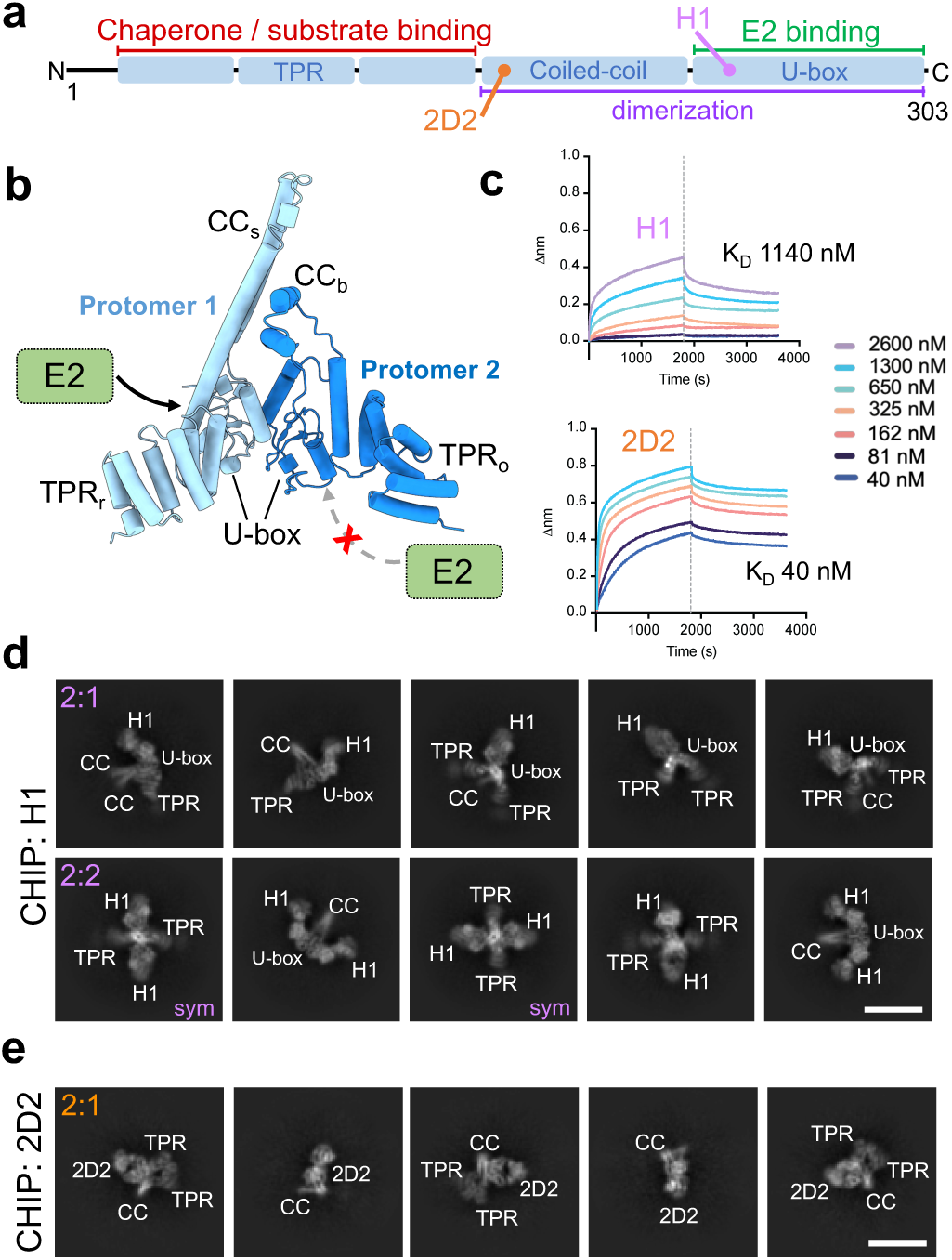
Selection of two Fabs that bind distinct sites on CHIP. (a) Schematic of human CHIP showing TPR domain (α1-6, residues 1-130), coiled-coil domain (α7-8, residues 131-224) and U-box domain (α8-10, residues 225-303) organization and functions. Approximate Fab (H1 and 2D2) binding sites marked as identified by cryo-EM (see below). (b) Crystal structure of murine CHIP adopts an asymmetric dimer conformation in which one protomer contains a “bent” coiled-coil domain (CC_b_) and a TPR domain positioned adjacent to the U-box in a manner that “occludes” E2 interactions (TPR_o_), while the other CHIP protomer contains a “straight” coiled-coil domain (CC_s_) and a TPR domain “rotated” away (TPR_r_), making U-box accessible for E2 binding (PDB: 2C2L)^13^ (c) BLI association and dissociation curves for CHIP binding to Fabs H1 and 2D2 shown as wavelength shift (delta nm) vs time (seconds). BLI was performed using biotinylated CHIP bound to streptavidin biological tips. K_D_ values (nM) derived from BLI are shown. Cryo-EM 2D class averages of (d) CHIP:H1 and (e) CHIP:2D2 showing 2:1 and 2:2 CHIP–Fab configurations with the CHIP TPR, U-box and CC domains and Fabs indicated (scale bar = 100 Å). Classes that were subsequently identified to represent symmetrical CHIP conformations (see below) are labeled “sym”.

Here, we sought to gain insight into CHIP mechanisms by determining structures using cryo-electron microscopy (cryo-EM). We took advantage of our recently isolated fragment antigen-binding (Fab) antibodies that target CHIP with a range of affinities and functional effects^27^. These Fabs were developed using the RAPID (Rare Antibody Phage Isolation and Discrimination) biopanning method^28^, with the goal of generating Fabs that stabilize the CHIP dimer structure and elucidate epitopes and regulatory mechanisms that might be therapeutically targeted^29,30^. Here we utilized two Fabs, H1 and 2D2, for CHIP structure determination, identified that they bind different epitopes, stabilizing the CHIP dimer in distinct states and with different functional outcomes. Structures of CHIP:H1 reveal Fab interactions with the CHIP U-box that overlap with the E2 binding site and stabilize three distinct configurations in which CHIP dimers adopt asymmetric and symmetric states through rearrangements of the CC and TPR domains. These conformational changes would enable ubiquitin-charged E2 (E2∼Ub) accessibility to both U-box sites and thus full activity of the CHIP dimer for substrate ubiquitination. In the CHIP:2D2 complex, we identify Fab binding to the asymmetric CHIP conformation, contacting both the bent and straight CC domains in a manner expected to block conformational changes required to form the symmetric state. In CHIP autoubiquitination assays H1 reduces ubiquitination, likely by inhibiting E2∼Ub binding, while 2D2 shows no substantial effect. Surprisingly, in our tau aggregation assays, we show that 2D2 reduces CHIP’s anti-aggregation activity, while H1 substantially increases this activity, suggesting that the H1-favored states enhance the anti-aggregation conformer. Based on these data, we propose models for CHIP function including roles for an asymmetric-to-symmetric conformational switch in its ubiquitination cycle, and how chaperone-like interactions of CHIP can support tau proteostasis.

## Results

### Two selected Fabs target distinct sites to stabilize CHIP dimers for cryo-EM

For cryo-EM structure determination of the CHIP dimer we sought to leverage the use of our recently-developed CHIP-targeting Fabs to improve stability for cryo-EM and overcome challenges based on CHIP’s small size (70 kDa as a dimer) and high flexibility^27,29,30^. We previously generated CHIP-Fabs using biopanning by phage display of a human naïve B-cell Fab library and screening methods: RAPID and BIAS (Biolayer Interferometry Antibody Screen)^27,28^. We first screened a set of CHIP-Fab candidates by cryo-EM 2D averaging analysis following incubations of 2 µM CHIP with 8 µM Fab and identified two Fabs with distinct complementarity-determining regions (CDRs), designated as “H1” and “2D2”, that improved stability and visualization of the CHIP dimer (Fig. 1c-e and Extended Data Fig. 1a). The binding affinities of these selected Fabs were determined using biolayer interferometry (BLI), showing that H1 binds CHIP with a modest, 1.1 µM affinity, while 2D2 binds with high affinity, at 40 nM (Fig. 1c and Extended Data Fig. 1b, c). In our initial attempts of CHIP incubations with selected Fabs H1 or 2D2, the cryo-EM 2D classification analysis could yield improved rigidity and visualization of Fab-bound CHIP, albeit with much preferred orientation, precluding further high-resolution 3D structure determination (Extended Data Fig. 2a). This issue of preferred orientation was overcome by the addition of Hsp70 protein, resulting in multiple orientations of well-resolved views of Fabs H1 or 2D2-stabilized CHIP (see Methods) (Fig. 1d, e). Subsequent 3D analysis of the complexes (see below) revealed no additional density corresponding to Hsp70, including its C-terminal EEVD residues that bind the TPR domain. Thus, we surmise that Hsp70 is unbound to the complex and potentially aids in cryo-protection, such as limiting CHIP interactions with the air-water interface^31^. We note that, in comparison, in the absence of the stabilizing Fabs, CHIP could not be resolved to high-resolution when characterized alone, or in the presence of Hsp70 protein or EEVD peptide (Extended Data Fig. 2b-d). Under the optimized conditions, our cryo-EM 2D classification analysis of Fab-stabilized CHIP revealed well-defined structural features of the complex in multiple orientations, including putative densities for the TPR, U-box and CC domains within CHIP dimers, along with densities for 1-2 bound Fabs (Fig. 1d, e). Monomeric CHIP was not observed in our analysis. Well-defined Fab densities indicated H1 and 2D2 bind at different sites on CHIP. H1 interacts with the U-box domains, while 2D2 contacts CC domains. Interestingly, we identified H1 in both 2:1 and 2:2 CHIP:Fab complexes, while 2D2 is only observed in a 2:1 complex. Overall, from these screening results we identify two Fabs H1 and 2D2 appear to target distinct sites and structural states of CHIP and enable, for the first time, cryo-EM structural characterization of the intact CHIP dimer.

### Structures identify Fab binding to U-box stabilizes asymmetric and symmetric CHIP dimer conformations

We next sought to determine 3D structures of CHIP bound to Fab H1. By 3D classification we identified 3 classes that exhibit distinct CHIP conformations and H1 binding stoichiometries: a predominant class (Class 1, with 50% of particles) with density for a single Fab bound to one side on the CHIP U-box dimer, and two additional classes (Class 2 with 21% and Class 3 with 29% of particles) containing two Fabs bound on either side of the U-box dimer (Fig. 2 and Extended Data Fig. 3a).

**Fig. 2.**
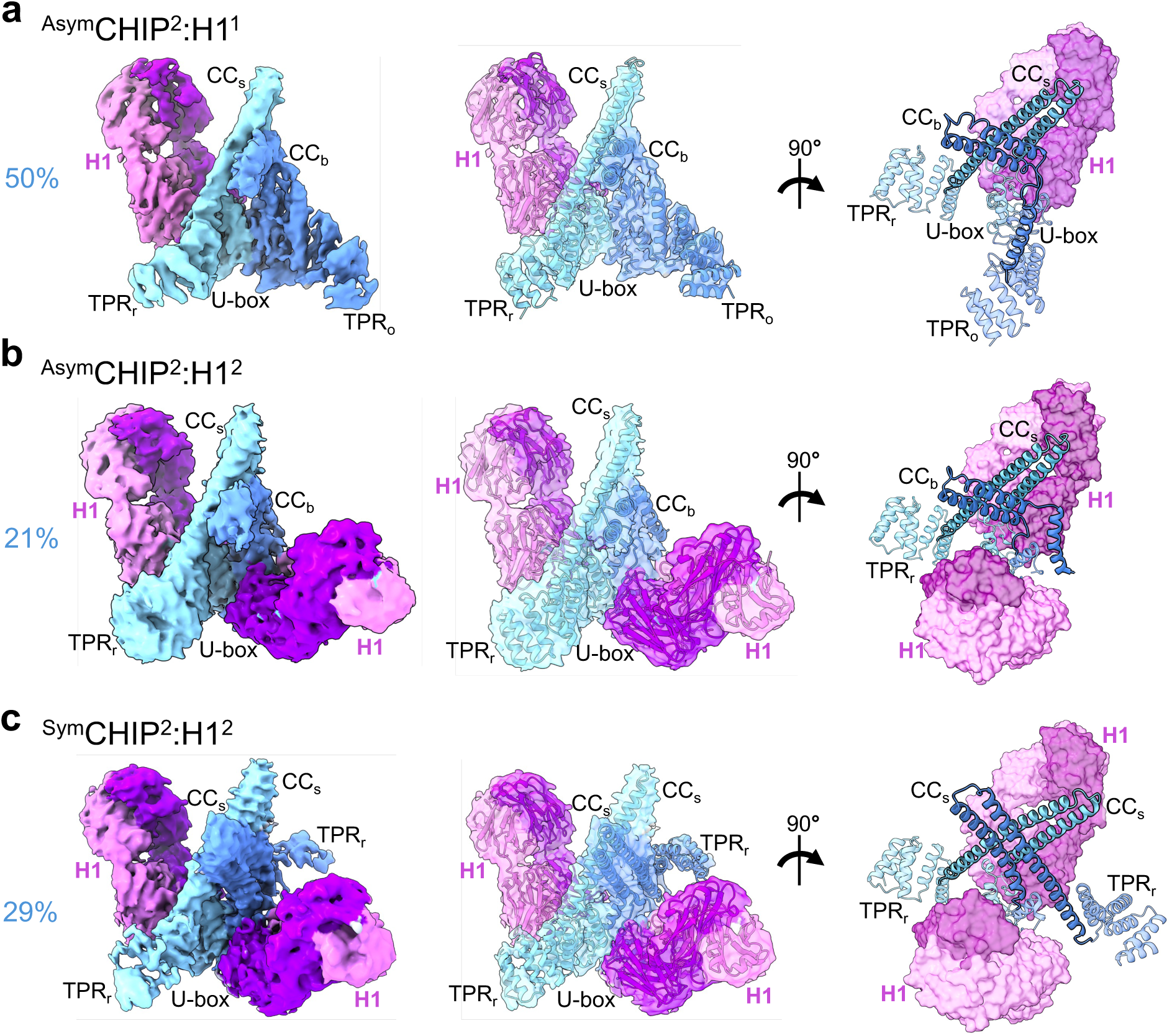
Cryo-EM map and models of CHIP:H1 identify three distinct conformations. Final cryo-EM density map (left), model fit in map (center) and rotated model view of the CHIP dimer U-box, TPR and CC domain positions (right) for: (a) ^Asym^CHIP^2^:H1^1^, showing the asymmetric CHIP dimer with straight (CC_s_) and bent (CC_b_) CC domains connected to respective rotated (TPR_r_) or occluding (TPR_o_) TPR domains with a single H1 Fab bound to a U-box domain (at 3.6 Å resolution), (b) ^Asym^CHIP^2^:H1^2^, showing the asymmetric CHIP dimer that resolves a single rotated TPR_r_ domain and two bound H1 Fabs bound to both sides of U-box dimer (at 3.9 Å resolution), and (c) ^Sym^CHIP^2^:H1^2^, showing CHIP in a symmetric conformation with two straight CC_s_ domains, two TPR_r_ domains, and two bound H1 Fabs bound to both sides of U-box dimer (at 3.7 Å resolution). CHIP is colored by protomer (light and dark blue) and Fab H1 is colored by heavy and light chains (magenta and pink, respectively).

Following refinement, Class 1 (denoted ^Asym^CHIP^2^:H1^1^) resolved to 3.6 Å overall resolution with density for H1 variable domains and CHIP U-box exhibiting the highest resolution (at 2-3.5 Å) while the TPR and CC domains are more flexible and at a lower (4-7 Å) resolution. (Fig. 2a and Extended Data Figs. 3a-c and 4a). An atomic model was achieved by rigid-body docking human CHIP and H1 Fab homology models based on crystal structures of murine CHIP (PDB: 2C2L) and a Fab (PDB: 1M71) and refinement using Rosetta (Extended Data Fig. 4b and Table 1). From the ^Asym^CHIP^2^:H1^1^ model we identify that CHIP adopts an asymmetric dimer conformation identical to previous crystal structure (Cα RMSD = 1.0 Å) (Fig. 2a and Extended Data Fig. 4i)^13^. While one CHIP protomer contains an intact straight CC domain and a rotated position of the TPR domain (TPR_r_), the opposing protomer contains a bent CC domain, forming a Y-shape dimer interaction with the straight CC, with the connected TPR domain in the occluding position (TPR_o_) (Fig. 2a). The straight CC domain contains a large, 52-residue helix α7 (residues F131-R182) extending from the TPR domain, forming an antiparallel hairpin with α8 (D190-K223) (Extended Data Fig. 4l). Conversely, in the opposing CC domain, helices α7 and α8 are shorter due to flexible strands between residues E152-S159 and V218-K223, respectively, resulting in a bent arrangement. In the CHIP protomer with bent CC domain, the unfolded portion of α7 enables the TPR_o_ domain to be positioned adjacent the U-box domain such that it occludes the E2 binding site, as previously described^13^ (Fig. 2a and Extended Data Fig. 4l). Notably, H1 interacts on the opposite dimer face, binding the U-box domain that is accessible for E2 binding resulting from the TPR_r_ position. This interaction is further characterized below. Importantly, these results identify that human full-length CHIP indeed adopts the unusual asymmetric dimer conformation in solution, without crystallographic constraints^13^. Moreover, the identification that a single Fab H1 binds to one U-box site, while the second site is unoccupied and occluded by TPR domain (Fig. 2a) supports the previously proposed mode of CHIP-E2 interactions for this conformation^13^.

**Table 1.** Cryo-EM data collection, refinement and validation statistics.

| Data collection and processing | AsymCHIP<sup>2</sup>:H1<sup>1</sup> | AsymCHIP<sup>2</sup>:H1<sup>2</sup> | SymCHIP<sup>2</sup>:H1<sup>2</sup> | CHIP U-box: H1 epitope | CHIP:2D2 complex | CHIP CC:2D2 epitope |
| --- | --- | --- | --- | --- | --- | --- |
| Microscope and camera | Titan Krios K3 | Titan Krios K3 | Titan Krios K3 | Titan Krios K3 | Titan Krios K3 | Titan Krios K3 |
| Magnification | 59,952 (K3) | 59,952 (K3) | 59,952 (K3) | 59,952 (K3) | 59,952 (K3) | 59,952 (K3) |
| Voltage (kV) | 300 | 300 | 300 | 300 | 300 | 300 |
| Data acquisition software | Serial EM | Serial EM | Serial EM | Serial EM | Serial EM | Serial EM |
| Exposure navigation | Image shift | Image shift | Image shift | Image shift | Image shift | Image shift |
| Electron exposure (e-/Å<sup>2</sup>) | 66 (K3) | 66 (K3) | 66 (K3) | 66 (K3) | 66 (K3) | 66 (K3) |
| Defocus range (μm) | -0.8 to -1.8 | -0.8 to -1.8 | -0.8 to -1.8 | -0.8 to -1.8 | -0.8 to -1.8 | -0.8 to -1.8 |
| Pixel size (Å) | 0.834 | 0.834 | 0.834 | 0.834 | 0.834 | 0.834 |
| Symmetry | C1 | C1 | C1 | C1 | C1 | C1 |
| Initial particle images (no.) | 1,082,688 | 1,082,688 | 1,082,688 | 1,082,688 | 962,842 | 962,842 |
| Final particle images (no.) | 482143 | 202245 | 278356 | 482143 | 85066 | 337834 |
| Map resolution (Å) | 3.6 | 3.9 | 3.7 | 3.1 | 4.8 | 3.4 |
| FSC Threshold | 0.143 | 0.143 | 0.143 | 0.143 | 0.143 | 0.143 |
| Map resolution range (Å) | 2-7 | 2-7 | 2-7 | 2-7 | 3-9 | 2.4-7 |
| **Refinement** |  |  |  |  |  |  |
| Model resolution (Å) | 3.6 | 4.0 | 3.8 | 3.5 | 4.9 | 3.5 |
| FSC threshold | 0.143 | 0.143 | 0.143 | 0.143 | 0.143 | 0.143 |
| Map sharpening B factor (Å<sup>2</sup>) | -176.5 | -161.8 | -165.7 | -108.6 | -247.9 | -175.5 |
| **Model Composition** |  |  |  |  |  |  |
| Non-hydrogen atoms | 7818 | 5306 | 5742 | 2310 | 3972 | 4360 |
| Protein residues | 996 | 1325 | 1434 | 303 | 992 | 562 |
| Ligands | 0 | 0 | 0 | 0 | 0 | 0 |
| **R.m.s deviations** |  |  |  |  |  |  |
| Bond lengths (Å) | 0.012 | 0.009 | 0.009 | 0.011 | 0.009 | 0.012 |
| Bond angles (degrees) | 1.632 | 1.739 | 1.677 | 1.418 | 1.647 | 1.908 |
| **Validation** |  |  |  |  |  |  |
| MolProbity score | 0.84 | 0.82 | 0.67 | 0.89 | 0.86 | 1.00 |
| Clashscore | 0.06 | 0.00 | 0.00 | 0.00 | 0.00 | 0.12 |
| Rotamer outliers (%) | 0.00 | 0.00 | 0.00 | 0.00 | 0.00 | 0.00 |
| **Ramachandran plot** |  |  |  |  |  |  |
| Favored (%) | 95.55 | 95.35 | 97.05 | 94.28 | 94.82 | 92.96 |
| Allowed (%) | 3.85 | 4.04 | 2.74 | 5.39 | 4.67 | 6.50 |
| Disallowed (%) | 0.61 | 0.61 | 0.21 | 0.34 | 0.51 | 0.54 |

Refinement of Class 2 (denoted ^Asym^CHIP^2^:H1^2^) resulted in an overall map resolution of 3.9 Å and shows two H1 molecules bound on either side of the U-box dimer (Fig. 2b and Extended Data Figs. 3d, e and 4c). Density for only a single TPR domain was observed. Following molecular modeling as above, we identify CHIP remains in the asymmetric dimer conformation with straight and bent central CC domains, as seen in Class 1 (Fig. 2b and Extended Data Fig. 4d and Table 1). The resolvable TPR domain with density is connected to the straight CC, again in the same rotated position as in Class 1. However, unlike Class 1, the presence of a second H1 appears to have displaced the second TPR domain connected to the bent CC domain (Fig. 2b and Extended Data Fig. 4j). The bent CC domain has a partially resolved helix α7 (residues A138-I157) that is shifted by ∼45° compared to Class 1, indicating that the connected TPR domain is in a different position (Extended Data Fig. 4k-l). The two H1 molecules are oriented symmetrically and bind the same U-box sites on either side of the dimer (Fig. 2b). Overall, the Class 2 ^Asym^CHIP^2^:H1^2^ structure captures a novel state of CHIP dimer in which both E2 binding sites on U-boxes are accessible. The second site interaction displaces the TPR from U-box, resulting in loss of its density. This likely occurs because the TPR is flexibly connected to the bent CC domain through the unfolded N-terminal portion of helix α7 (unresolved residues F131-S137), compared to the adjacent protomer where straight CC helix α7 is intact and rigidly extends from the TPR_r_ domain.

Class 3 (denoted ^Sym^CHIP^2^:H1^2^) refined to 3.7 Å overall resolution and also contains two H1 molecules bound on either side of U-box dimer (Fig. 2c, and Extended Data Figs. 3f, g and 4e). Density for both TPR domains is identified in an apparent symmetric orientation but is more poorly resolved for one CHIP protomer, limiting modeling accuracy. Intact densities for the CC domain helices α7-8 are observed for both CHIP protomers in addition to well-defined density for both H1s and the U-box dimer. Strikingly, molecular modeling reveals CHIP and H1 are in a symmetric arrangement with both CC domains adopting the straight conformation in which helix α7 (residues F131-R182) is intact and forms an antiparallel helical hairpin with α8 (Fig. 2c and Extended Data Fig. 4f, k, l and Table 1). Thus, the CC domains are in a distinct symmetric cross arrangement not observed in our other CHIP:H1 structures or the previous intact full-length crystal structure^13^. We note this CC arrangement is similar to a previous truncated CC-U-box dimer structure^14^. Additionally, the TPR domains are in a symmetric orientation, both rotated away from U-box, enabling H1 access to both U-box sites (Fig. 2c). Overall, Class 3 ^Sym^CHIP^2^:H1^2^ structure captures a distinct symmetrical CHIP dimer state in which both CC domains adopt the straight conformation with intact α7 helices that extend from TPR domains, stabilizing both TPR orientations rotated away from U-box domains.

### CHIP-H1 interactions mimic U-box binding by cognate E2 enzymes

Our structures of CHIP:H1 reveal the Fab interaction site appears similar to where E2 ubiquitin conjugating enzymes contact U-box domains during Ub transfer^25,32^. To further define the epitope, we performed focused refinement using a mask including the U-box and H1 heavy and light chain variable domains (V_H_ and V_L_) (Extended Data Fig. 3a). A focused map at 3.1 Å was determined that resolves many side chain densities, enabling accurate modeling of the U-box:H1 interface (Fig. 3a and Extended Data Figs. 3h, i and 4g, h and Table 1). Based on the model we identify H1 interacts with the CHIP U-box domain through extensive hydrophobic and polar contacts (largely through H1 V_L_) that bury a surface area of *∼*1027 Å^2^ (Fig. 3a, b). H1 V_L_ CDR3 loop residues (102-RSGYSGYDNNLGAFD-116) make hydrophobic contacts with a conserved hydrophobic groove on the U-box surface comprised of loops α8-9 (229-239) and α9-10 (264–275), and helix α9 (254–264) (Fig. 3b). The V_L_ CDR3 interaction is also supported by apparent salt bridges between CDR3 G107 backbone carbonyl and CHIP R272, and between CDR3 D116 and CHIP S236 backbone carbonyl. CHIP F237 packs between H1 V_L_ CDR3 R102 and D116 sidechains forming another hydrophobic interaction. Side chains of H1 V_L_ CDR1 Y36 and CDR2 Y58 also make hydrophobic contacts with CHIP K234, F237 and CHIP T271 respectively. The H1 heavy chain V_H_ CDR loops form a network of polar contacts with CHIP, including hydrogen bonds (CDR2 S72–CHIP E238, and CDR1 Y48–CHIP R263), and a salt bridge between CDR2 K69 and CHIP E259, along with hydrophobic contacts of CDR2 Y65 and CHIP I235.

**Fig. 3.**
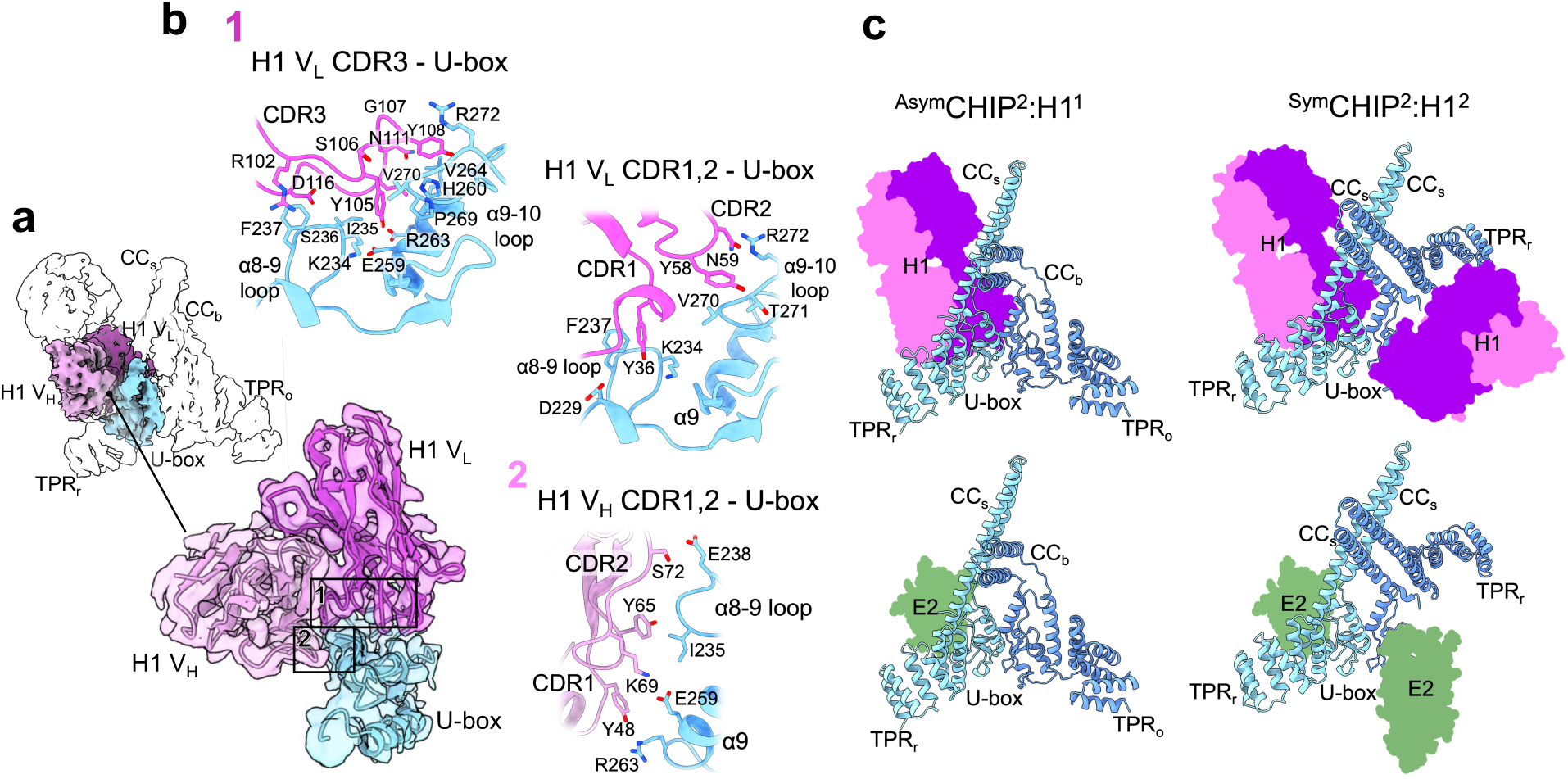
Fab H1 CDR interactions with the CHIP U-box and comparison to E2 binding. (a) CHIP:H1 interaction epitope focused refinement map at ∼3.1 Å resolution (shown overlaid on transparent ^Asym^CHIP^2^:H1^1^ map) with enlarged view of model fit in map showing H1 variable light (V_L_, magenta) and heavy (V_H_, pink) chain CDR contacts with the CHIP U-box (light blue), highlighting 2 distinct CDR contact regions (boxed). (b) Enlarged views of the H1 CDR interactions with CHIP U-box domain showing contacts between H1 V_L_ CDRs 1,2,3 and U-box α8-9 and 9-10 loops and α9 residues, and between H1 V_H_ CDRs 1,2 and U-box α8-9 loop and α9 residues. (c) Models of ^Asym^CHIP^2^:H1^1^ (left) and ^Sym^CHIP^2^:H1^2^ (right) with single and double H1 Fab molecules (colored as above) bound to the U-box sites compared to models of CHIP in the respective asymmetric (lower, left) and symmetric (lower, right) conformations with single and double E2 Ubc13 molecules (green) binding to the same U-box sites, based on the CHIP U-box:Ubc13 complex crystal structure (PDB: 2C2V)^13^.

Based on structural comparisons, we identify that the residues within the CHIP U-box hydrophobic groove that interact with Fab H1, are strikingly similar to those involved in CHIP-E2 interactions, as observed in previous U-box:E2 crystal structures including CHIP:Ubc13 and CHIP:UbcH5^13,14^ (Extended Data Fig. 5). Indeed, alignment of U-box:Ubc13 dimer (PDB: 2C2V) onto our cryo-EM models for ^Asym^CHIP^2^:H1^1^ and ^Sym^CHIP^2^:H1^2^ CHIP dimer states show substantial overlap and similar binding modes of H1 and E2 with CHIP (Fig. 3c). Notably, CHIP residues H260, I235, R272, and V264, which contact the conserved “S-P-A motif” in loop 7 of cognate CHIP-binding E2s^32^ interact with H1 V_L_ CDR3 residues Y105, S106, G107 and Y108 (Extended Data Fig. 5). Additionally, while H1 binds CHIP with a modest 1.1 µM affinity (see Fig. 1c), which is among the weakest contacts observed for the Fabs generated^27^, E2 interactions with CHIP are similarly weak (reported ∼1-10 µM for Ubc13^13^ and UbcH5A^14^, and much weaker for other E2-U-box/RING domain interactions^33^). Thus, we observe striking structural and functional similarities between interactions by Fab H1 and cognate E2s with CHIP U-box.

The ^Sym^CHIP^2^:H1^2^ structure also reveals how the symmetric position of rotated TPR_r_ domains would allow two E2s to bind CHIP dimers simultaneously, as we identified for H1, without any structural hinderance (Fig. 3c). However, it remains unclear how a ubiquitin-charged E2 (E2∼Ub) would interact with CHIP dimers. To further explore this, we generated models of 2:2 CHIP:E2-Ub complexes (with CHIP-binding E2s Ubc13, UbcH5A or Ube2w)^13–15^ using AlphaFold 3^34^ (Extended Data Fig. 6). Strikingly, the predicted structural models consistently show symmetrical CHIP dimers that are identical to ^Sym^CHIP^2^:H1^2^ containing two straight CC domains and rotated TPR_r_ domains, enabling simultaneous E2-Ub binding to both U-box sites. Additionally, the AlphaFold 3 models also predict specific *trans-* interactions of the E2-bound Ubs with the U-box and TPR domains of the opposite symmetrically positioned CHIP protomers, which, to our knowledge, have not been described before, and may be important in mediating Ub transfer (Extended Data Fig. 6). Together, these predicted structures further support a role for the symmetric conformation of CHIP dimers observed in ^Sym^CHIP^2^:H1^2^, in the E2 binding interactions and ubiquitination.

### Fab 2D2 binds CHIP across the straight and bent CC domains at the dimer interface

We next sought to determine structures of CHIP bound to Fab 2D2 by cryo-EM. As discussed above, unlike for H1, only a 2:1 CHIP:2D2 complex is observed in 2D class averages (see Fig. 1e). For 3D classification and refinement, only particles containing density for the full CHIP dimer were initially selected based on 2D analysis (Extended Data Fig. 7a). The final map for CHIP:2D2 refined to an overall 4.8 Å resolution (Fig. 4a and Extended Data Fig. 7a-c, and Table 1). We contend the lower resolution compared to CHIP:H1 is likely due to Fab binding to the more flexible CC domains compared to the U-box. The CHIP dimer and 2D2 homology models were rigid-body docked into the map (Fig. 4a) and indicate that CHIP adopts an asymmetric dimer conformation that is similar to ^Asym^CHIP^2^:H1^1^ and the previous crystal structure^13^. Notably, 2D2 appears to contact the CHIP straight and bent CC domains dimerization interface, and thus likely stabilizes the asymmetric CHIP dimer conformation.

**Fig. 4.**
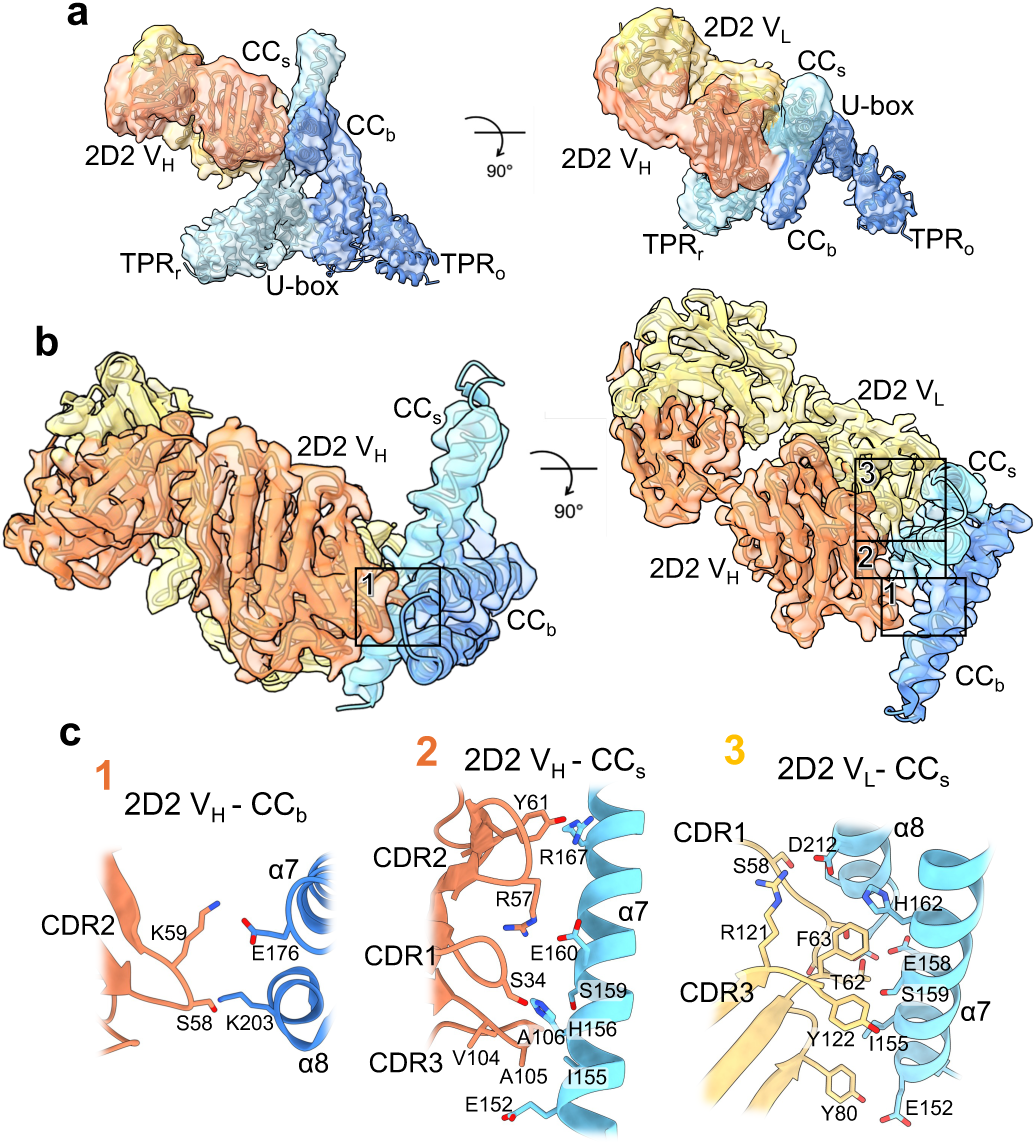
Cryo-EM structures of CHIP bound to Fab 2D2 identify interactions with the asymmetric CC interface. (a) Views of CHIP:2D2 cryo-EM density map at 4.8 Å resolution and rigid-body docked model, colored to indicate Fab 2D2 heavy (V_H_, orange) and light (V_L_, yellow) chains and CHIP protomers (colored as above). (b) Focused, refined map and model of the 2D2 interaction with the CHIP CC domains at 3.4 Å resolution, highlighting 3 distinct CDR contact regions (boxed). (c) Enlarged views of the 2D2 CDR interactions with CHIP CC domains showing contacting residues between the 2D2 V_H_ CDRs and CHIP CC_b_ α7-8 (left) and CC_s_ α7 (middle), and between the 2D2 V_L_ CDRs and the CHIP CC_s_ α7-8 (right).

To obtain details of the 2D2-CHIP interaction, focus refinement was performed using a mask around 2D2 and CHIP CC domains and the full particle set. The map improved to ∼3.4 Å resolution and exhibited well-resolved sidechain densities at the 2D2-CHIP CC interface (Fig. 4 b, c and Extended Data Fig. 7a, d-f and Table 1). The refined model reveals that 2D2 indeed engages the CC domains across both CHIP protomers, interacting with residues from both helices α7-8. All 2D2 V_H_ and V_L_ CDR loops engage at specific sites on CC domains, encompassing a buried surface area of ∼1070 Å^2^. Most notably, V_H_ CDR2 (residues: 57-RSKWY-61) makes multiple contacts with both the bent (CC_b_) and straight (CC_s_) domains, forming salt bridge interactions between K59 and CC_b_ E176 and between R57 and CC_s_ E160, while polar contacts are observed between S58 and CC_b_ K203, and between Y61 and CC_s_ R167 (Fig. 4c). Additionally, V_H_ CDR1 S34 hydrogen bonds with CC_s_ H156, while residues 104-VAA-106 in V_H_ CDR3 make hydrophobic interactions with residues CC_s_ I155, H156 and S159. Conversely, V_L_ CDRs (including CDR1 residues 58-SVSSTF-63) interact primarily with the protomer containing CC_s_, contacting hydrophobic and polar residues (I155, E158, H162, S216, D219) along an exposed helical face of α7,8, while a salt bridge contact is also identified between V_L_ CDR3 R121 and D212 in α8 of CC_s_.

In summary we identify that 2D2 makes extensive hydrophobic and polar interactions with CHIP along helices α7-8 of both the bent and straight CC domains, supporting the observed high-affinity of 2D2-CHIP binding (see Fig. 1c). Further, when we overlay 2D2 onto the symmetric CC dimer conformation from the ^Sym^CHIP^2^:H1^2^ structure, we identify that the shifted position of the second straight CC would be incompatible for CHIP-2D2 contacts (Extended Data Fig. 7g). Indeed, 2D2 contacts show that it specifically recognizes the asymmetric CHIP dimer state which has a single U-box site for E2 binding, further supporting that this is the predominant solution state. Additionally, this interaction likely stabilizes the CHIP asymmetric state by preventing conformational changes of the CC dimer and connected TPR domain that we identify with H1.

### Interactions by Fabs H1 and 2D2 alter CHIP function in ubiquitination and tau anti-aggregation

Based on the different Fab H1 and 2D2 interaction sites identified in our cryo-EM structures, we postulated that their binding may alter distinct aspects of CHIP function. We first investigated CHIP autoubiquitination in the presence of H1 or 2D2 using UbcH5b (also known as UBE2D2), an established E2 in the CHIP ubiquitination cycle^14^. In the absence of Fabs we observe robust poly-ubiquitination of CHIP, as indicated by the ladder of high molecular weight bands by Western, when blotted for CHIP (Fig. 5a). In the presence of increasing amounts of H1 (2-8 µM, compared to 2 µM of CHIP) under the same conditions, CHIP autoubiquitination is substantially reduced, with a near complete loss at 8 µM H1. This demonstrates H1 indeed inhibits CHIP-mediated ubiquitination, as predicted by our CHIP:H1 structures. Thus, under these conditions H1 likely outcompetes E2∼Ub binding to the U-box, inhibiting Ub transfer. The high concentration of H1 required for complete inhibition is expected given the 1.1 µM affinity determined for H1, which is equivalent to reported ∼1-10 µM (or weaker) affinities of E2s for U-box/RING E3s^13,14,33^. Conversely, under increasing concentrations of 2D2 (1-6 µM) we observe no change in CHIP-autoubiquitination, indicating 2D2 binding does not affect E2∼Ub binding and Ub transfer (Fig. 5a), as predicted by our cryo-EM structures, which identify 2D2 binds to the CC domain dimer in the asymmetric state, away from the E2 binding site (see Fig. 4).

**Fig. 5.**
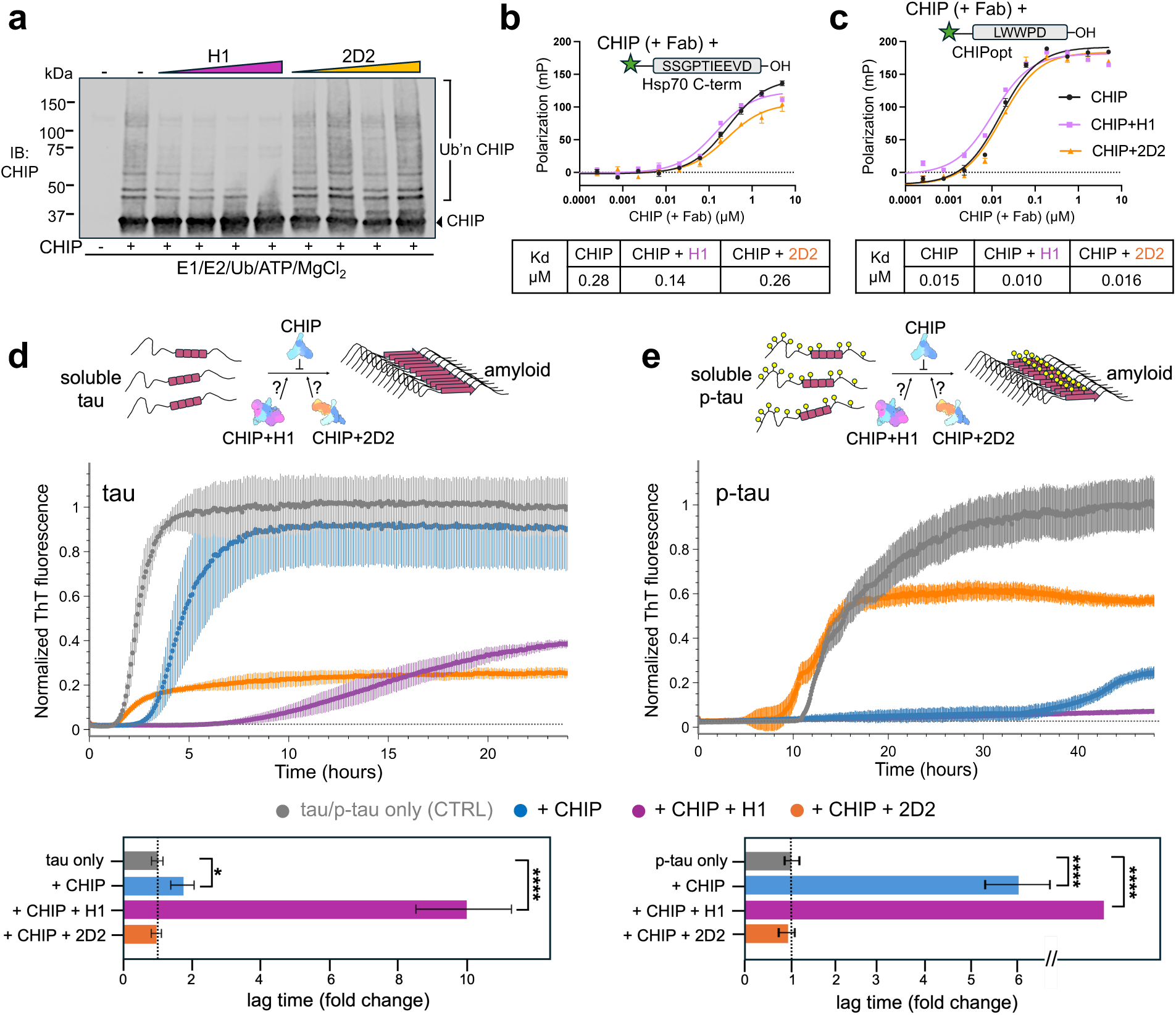
Fab interactions alter ubiquitination and tau anti-aggregation functions of CHIP. (a) *In vitro* autoubiquitination of CHIP (2 μM) in the presence of increasing amounts of Fabs H1 (2,4,6,8 μM) and 2D2 (1,2,4,6 μM). The reactions were incubated for 15 mins at room temperature, quenched with SDS–PAGE loading buffer, and analyzed by western blotting (IB:CHIP). Assay was performed once. (B-C) FP-based CHIP-TPR binding assay measuring the binding of (b) FAM-labeled Hsp70 C-terminal peptide (5FAM-Ahx-SSGPTIEEVD) or the binding of (c) a higher-affinity CHIP-optimized peptide “CHIPopt” FITC-Ahx-LWWPD^22^ to CHIP in the presence of H1 and 2D2. Binding curves were fit using non-linear regression and Kd values (µM) for peptide binding for CHIP alone, CHIP with H1, or CHIP with 2D2 are shown below. Experiments were performed in technical triplicates. (D-E) Tau aggregation schematic and normalized ThT fluorescence to detect (d) unmodified 0N4R tau or (e) phosphorylated 0N4R tau (p-tau) aggregation over time (hours) either alone (grey), or in the presence of CHIP (blue), CHIP with H1 (magenta), or CHIP with 2D2 (orange). Data is shown as mean ± SD (n=3) and incubations were with equimolar (5 μM) concentrations of tau (or p-tau), CHIP, and Fabs (H1 or 2D2). Derived lag time changes for each condition are shown below as the fold change relative to unmodified tau and p-tau aggregation alone. Data is normalized to free tau and represented as mean ± SD. Statistical significance was determined by one-way analysis of variance (ANOVA) with Dunnett’s post-hoc test (*p<0.01,****p<0.0001).

We next tested peptide binding to CHIP TPR domain to assess whether the conformational changes we observe in CHIP dimers impact TPR function. Using our established FP peptide binding assay^35^ we measured the binding affinity of Hsp70 C-terminal EEVD peptide (5FAM-Ahx-SSGPTIEEVD) to CHIP in the presence of H1 or 2D2 (Fig. 5b). In agreement with previous studies, we determined an apparent Kd of 0.28 µM between CHIP and the EEVD peptide alone^35^. In the presence of equivalent amounts of H1 or 2D2 we determined apparent affinities of 0.14 µM and 0.26 µM, respectively, indicating no substantial reduction in EEVD-TPR binding affinity. The same tests were conducted using an optimized Hsp70-EEVD peptide (“CHIPopt” FITC-Ahx-LWWPD) that binds CHIP TPR with a ∼10-fold tighter affinity^22^ (Fig. 5c). With CHIPopt we determined apparent Kds of 0.015 nM for CHIP, compared to 0.010 nM and 0.016 nM when H1 or 2D2 are included, respectively. Thus, TPR domains remain competent for binding in the presence of either Fab H1 and 2D2. The consistent ∼2-fold increase in affinity observed in the presence of H1 is intriguing and suggests that H1 may partially enhance TPR binding to EEVD substrates.

Finally, we sought to explore whether H1 and 2D2 impact CHIP’s ability to block aggregation of tau protein *in vitro*. As noted above, in recent studies we discovered that CHIP has chaperone-like or “holdase” functions as a potent inhibitor of tau fibrilization, attributed to direct binding to soluble tau through the TPR domain^12^. We identified the presence of CHIP results in a ∼2-fold increase in the delay of tau aggregation onset *in vitro* (termed “lag time”) for unmodified 0N4R tau, and a much greater ∼5-fold increase in lag time for the disease-relevant phosphorylated 0N4R tau (p-tau)^12^’^36–38^. Here, using our established assay measuring heparin-induced tau aggregation by thioflavin T (ThT) fluorescence^12^, we questioned whether H1 or 2D2 interactions with CHIP modulate its anti-aggregation activity. For unmodified tau, we identify the presence of H1 increased the aggregation lag time substantially by 10-fold compared to tau alone (i.e. 5-fold above that for tau incubated with CHIP alone) (Fig. 5d). Conversely, 2D2 abolished CHIP anti-aggregation effects; CHIP incubated with 2D2 exhibited no differences in lag time compared to unmodified tau alone. For p-tau fibrilization, the addition of H1 shows a striking enhancement of CHIP-mediated anti-aggregation, such that ThT fluorescence is not detected over the long experimental timeframe of 48 hours, indicating complete inhibition of tau fibrilization (Fig. 5e). 2D2 again abolishes CHIP’s anti-aggregation effect for p-tau. However, we consider this effect more substantial because CHIP alone has a much greater anti-aggregation effect on p-tau. We note the variability in fluorescence plateaus, and attribute this to potential heterogeneity in tau fibrilization or ThT binding and fluorescence properties, as previously reported^39^. In control experiments the presence of H1 or 2D2 alone, without CHIP, did not alter fibrilization of tau/p-tau (Extended Data Fig. 8). Together, these results show that H1 and 2D2 binding to CHIP have significant and opposing effects on CHIP’s chaperone-like anti-aggregation activity on both tau and p-tau.

Based on the epitope sites identified in our structures this indicates that Fab binding to U-box greatly enhances CHIP’s anti-aggregation activity, while Fab binding to CC dimer abolishes this effect. Although the specific anti-aggregation mechanism is unclear, we expect Fab binding likely causes direct or indirect changes that alter CHIP binding interactions with soluble tau. Considering previous work identifying tau interacts with the TPR and CC domains^12,24^, changes in accessibility to these sites likely contributes to the tau anti-aggregation effects observed. Thus, H1’s potent enhancement may be through stabilization of CHIP dimer states that favor TPR interaction with tau, such as the symmetric state in which both TPRs are rotated away from U-boxes. Conversely, 2D2-CC dimer contacts may directly block interactions between tau and CC domain^24^ or limit conformational states, such as the symmetric dimer, that may favor tau binding. Importantly, these results highlight the substantial modulating effects Fab interactions have on CHIP functions, revealing how 2D2 and H1 could be used to partition CHIP towards ubiquitination or chaperone-like/holdase pathways.

## Discussion

Ubiquitination and degradation are essential multistep protein quality control processes required for protein turnover and clearance. The U-box E3 ligase CHIP functions as a critical triage hub for many pathological proteins, coordinating Ub transfer to substrates delivered through Hsp chaperone-dependent and independent pathways^1,3,10,12,22,23^. Moreover, CHIP’s diverse functions may extend to chaperone-like/holdase activity that slows tau aggregation^12^. The structural basis for these diverse functions has remained elusive.

To overcome challenges in structural characterization of CHIP, we utilized recombinant Fabs identified by an antibody biopanning campaign^27^. By cryo-EM, we identify that Fab H1 binds the CHIP U-box domain, contacting sites that mimic cognate E2 interactions^32^ and stabilizing distinct asymmetric and symmetric conformational states of CHIP dimer (Figs. 2, 3). This interaction limits CHIP autoubiquitination, likely by inhibiting E2 binding, but dramatically improves CHIP-dependent inhibition of tau aggregation (Fig. 5). Fab 2D2 binds at the coiled-coil interface of asymmetric CHIP dimer, interacting with both straight and bent conformations of helices α7-8, likely blocking conformational changes to the symmetric state identified in our structures with H1 (Fig. 4). 2D2 binding does not alter CHIP autoubiquitination but does abolish the inhibition of tau aggregation by CHIP (Fig. 5). Together these results uncover previously unknown states of the intact CHIP dimer and suggest how these states and interaction surfaces contribute to distinct functions.

Based on our cryo-EM structures and functional analysis, we propose models for the CHIP ubiquitination cycle and chaperone-independent interactions with tau (Fig. 6). The three structures of CHIP:H1 suggest that CHIP undergoes an asymmetric-to-symmetric conformational switch involving rotation of TPR domain and extension of the bent CC domain (Fig. 6a and Supplementary Video1). In this model, the symmetric dimer conformation allows E2∼Ub accessibility to both U-box sites, while the asymmetric state is “half-of-sites” active (Fig. 6a). We propose the ^Asym^CHIP^2^:H1^2^ conformation in which the displaced TPR is flexible and not resolved is intermediate between these states. Thus, TPR rotation would potentially occur first, releasing constraints on bent CC and enabling helix α7 to form an extended orientation and adopt the straight conformation seen in the symmetric state. As a result of these conformational changes, the U-box sites would flip between accessible and occluded states across protomers, explaining how both sites are active during ubiquitination but why only one E2∼Ub may be able to bind at any given time, as previously proposed (Supplementary Video1). Fab 2D2 binding to the CC domains of the asymmetric dimer prevents this conformational switch, but it retains E2∼Ub accessibility to U-box of one protomer with the rotated TPR domain enabling ubiquitination to proceed normally. From our ^Sym^CHIP^2^:H1^2^ structure and AlphaFold 3 analysis (Fig. 2 and Extended Data Fig. 6), we propose that the symmetric CHIP dimer is compatible for binding by two E2∼Ub complexes, forming a 2:2 complex that would be a full-site active state for Ub transfer. The symmetric state may be more transient and specific to H1 binding considering it is less populated in our cryo-EM data (29% compared to 50% for the asymmetric state) and not previously observed crystallographically as a full-length dimer. However, in support of an active state symmetric dimer, a truncated CHIP dimer crystal structure without TPR domains resolves a symmetric CC interaction^14^. Additionally, MD simulations of CHIP indicate the bent and straight CC conformations interconvert and may assemble initially to form a symmetric dimer^26^. Considering H1 interactions with CHIP show remarkable similarity to E2 contacts, a compelling model supported by our structures is that E2∼Ub binding to the accessible U-box triggers asymmetric-to-symmetric conformational changes in CHIP that unleash the second U-box for a second E2∼Ub interaction. Thus, the conformational switch may be kinetically triggered (by H1 or E2∼Ub binding), as previously proposed^26,40^, which could regulate ubiquitination of certain substrates or facilitate E2 coordination for Ub chain elongation steps^41^. Additional structures of intact CHIP dimer-E2∼Ub complexes are needed to further define CHIP’s active state(s).

**Fig. 6.**
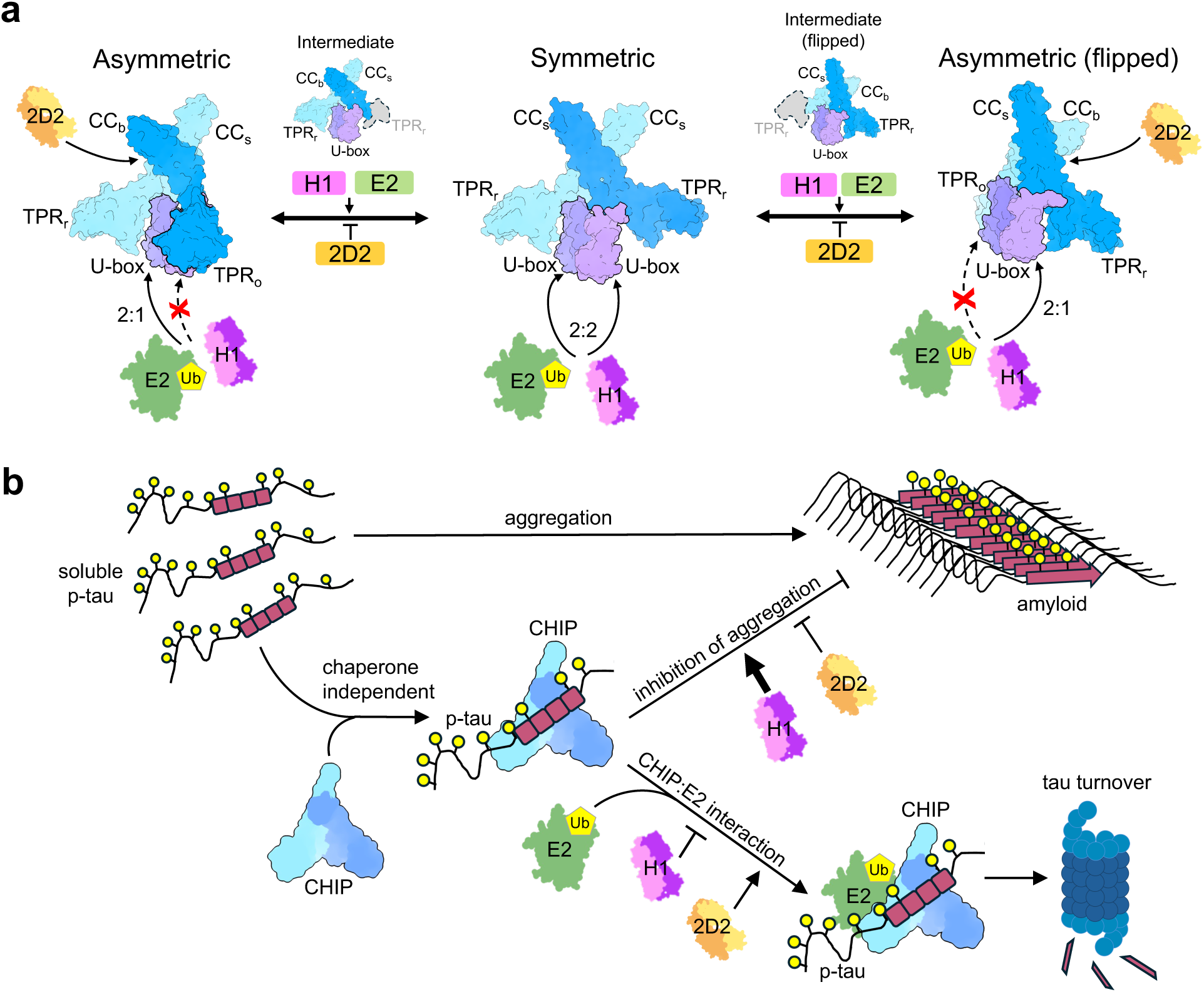
Models for CHIP conformational cycle and function during ubiquitination and tau anti-aggregation. (a) A structural schematic of CHIP conformational changes during E2-Ub binding and ubiquitination based on CHIP:H1 and CHIP:2D2 structures. CHIP primarily adopts the “Asymmetric” state in which the U-box site of one protomer is accessible to E2 (or Fab H1) binding due to the rotated TPR_r_ domain position while the U-box of the second protomer is occluded by its TPR_o_ domain. Conformational changes in CHIP (involving CC_b_ to CC_s_ and TPR_o_ to TPR_r_ domain changes in one protomer) result in a “Symmetric” state in which both U-box domains are accessible for E2 (or H1) binding, enabling formation of a 2:2 tetramer complex. This occurs through an asymmetric “Intermediate” in which the TPR_o_ is first displaced from the U-box interaction while the CC domains remain in the straight (CC_s_) and bent (CC_b_) configuration. These conformations enable the protomers flip between states, thus enabling the U-box sites to be equally accessible. CHIP protomers are colored as light and dark blue, with corresponding U-box domains colored purple and violet, respectively. (b) Model depicting CHIP function in binding soluble p-tau through its TPR and CC domains, preventing aggregation and amyloid formation and promoting chaperone-independent ubiquitination and turnover. Fab H1 binding strongly enhances the inhibition of tau aggregation by CHIP, while 2D2 binding abolishes this effect. These opposing functions are likely driven by distinct Fab interactions: H1 binds to the U-box domain, while 2D2 targets the CC domain, leading to enhanced or inhibited CHIP–tau interactions, respectively.

Chaperone-independent functions of CHIP include ubiquitination of unmodified, phosphorylated or caspase-cleaved proteoforms of tau in the absence of chaperones^12,22,24^, via direct contacts between CHIP TPR and CC domains and tau repeat domain^24^. CHIP also inhibits tau aggregation through TPR domain contacts^12^. This indicates dual chaperone-independent pathways for CHIP’s role in tau proteostasis: ubiquitination activity through E2∼Ub recruitment and “holdase” functions to prevent tau accumulation (Fig. 6b). From our structures and anti-aggregation analysis we identify H1 binding to U-box and 2D2 binding to CC sites, results in drastic differences in the anti-aggregation of p-tau (Fig. 5e). For H1, we propose interactions at U-box enhance CHIP contacts with tau which, thereby potentiate the anti-aggregation activity (Fig. 6b). This could occur through promotion of the asymmetric-to-symmetric conformational switch, which likely exposes additional interaction surfaces at sites of rearrangement in TPR and CC domains. While H1 interactions block E2∼Ub binding and likely limit tau ubiquitination, given the binding similarities between H1 and E2s, we postulate E2∼Ub binding to CHIP on its own may enhance CHIP-tau interactions for ubiquitination and proteasomal degradation in the chaperone-independent pathway. Conversely, 2D2 likely blocks a tau interaction site in CC domain or inhibits conformational changes that promote tau binding, resulting in a loss of the anti-aggregation activity. Importantly, our results reveal specific sites on CHIP that can be targeted by single-chain variable fragment (scFv) versions of the Fabs, or potentially peptides or small molecules, for modulating distinct CHIP functions *in vivo*.

Development of tools that can modulate CHIP’s activities have largely remained underexplored^27,42^. One of the goals of our study was to use Fabs to reveal possible protein surfaces on CHIP that might be targeted to tune its various functions as putative therapeutics. Such an approach has been successful in other systems *in vivo*, where modulation of molecular pathways has been achieved using Fab-derived therapeutics^29,43,44^. Here, we anticipate that the observed H1-mediated enhancement of CHIP’s anti-aggregation activity, likely mediated through promotion of conformational changes in CHIP, might guide the development of a CHIP activating tool.

## Methods

### Protein purification

WT CHIP (human, His-tagged) was expressed from a pET151 construct containing an N-terminal, tobacco etch virus (TEV)-cleavable 6xHis tag. *E. coli* culture was grown in LB broth at 37 °C, induced with 1 mM IPTG in log phase, cooled to 16 °C, and grown overnight. Cells were harvested, resuspended in binding buffer (40 mM Tris pH 8.0, 10 mM imidazole, 500 mM KCl) supplemented with protease inhibitors, and lysed via sonication. The clarified lysate was applied to Ni-NTA resin (Novagen), followed by washing with binding buffer and His wash buffer (40 mM Tris pH 8.0, 30 mM imidazole, 100 mM KCl). The protein was eluted using His elution buffer (40 mM Tris pH 8.0, 300 mM imidazole, 100 mM KCl). To remove the N-terminal His tag, proteins were dialyzed overnight at 4 °C with TEV protease. The digested material was then reapplied to Ni-NTA resin to separate the cleaved His tag, any undigested protein, and TEV protease. Finally, the protein was further purified using size exclusion chromatography (SEC) on a Superdex 200 column equilibrated in 20 mM HEPES, 100 mM KCl, pH 7.5. Aliquots were flash-frozen in liquid nitrogen and stored at −80°C for future use. WT Hsp70 was purified as described previously^45^. Tau and p-tau were purified as described previously^12^.

### Biopanning

Fabs described in this study were isolated from a human naïve B-cell phage displayed Fab library (diversity 4.1 x10^10^)^46^ using the previously described biopanning method RAPID (Rare Antibody Phage Isolation and Discrimination)^28^. Briefly, biotinylated CHIP was immobilized to magnetic streptavidin beads (Dynabeads M-280 Streptavidin) and incubated with the phage library. Beads were washed to remove weak and nonspecific binders. Bound phage were amplified using M13K07 helper phage, and the resulting phage mixture was used as the starting library for the subsequent round of biopanning. Four total rounds of this process were performed with increasing stringency by reducing the CHIP to phage library ratio and increasing the number of washes. RAPID flow cytometry was employed to identify the most enriched round of binders. Output phage pools were labeled with NHS-FITC and incubated with CHIP-bound polystyrene beads (SPHERO^TM^ Streptavidin, SVP-30-5). Subsequently, bound phage were analyzed with flow cytometry (benchtop Beckman Cytoflex Analyzer) and fluorescent signals were measured and compared. Individual clones from Round 3 of biopanning were examined using BIAS (Biolayer Interferometry Antibody Screen)^28^, where periplasmic extracts of Fab expressions were analyzed using biolayer interferometry in a 96 well plate format. Multiple clones, including Fabs H1 and 2D2, were isolated that showed promising association and dissociation curves.

### Fab production and purification

Fabs H1 and 2D2 were prepared from fresh transformants of BL21(DE3) *E. coli* cells containing Fab expression plasmids. Single colony clones were picked and inoculated into 50 ml of 2xYT, 2% glucose, 100 mg/ml Ampicillin. Overnight starter cultures were incubated at 37°C with 200 rpm shaking. The starter cultures were then inoculated into 1 L of 2xYT, 0.1% glucose, 100 mg/ml Ampicillin to a final OD_600_ of 0.05. Protein expression was induced at OD_600_ = 0.6 with IPTG (1 mM final). Expressions were done overnight at 20°C with 200 rpm shaking. Periplasmic extracts were isolated using osmotic shock. Overnight cell cultures were harvested by centrifugation at 9000 g for 15 min followed by the addition of ∼15 ml of ice-cold TES buffer. The solution was resuspended thoroughly to achieve a homogenous mixture and incubated at 4°C for 1 hour with gentle shaking. 25 ml of ice-cold water was added, and the mixture was incubated further for 45 min. The periplasmic fraction (supernatant) was collected by centrifugation at 10,000 g for 30 min and loaded to Ni-NTA resin for affinity purification. Purified Fabs were dialyzed into 25 mM HEPES, 50 mM KCl (10 kDa MWCO) and SEC was performed to remove any impurities (Superdex 200 10/300GL column). Pooled SEC fractions from a single peak showed only bands representing Fab on SDS-PAGE.

### Biolayer Interferometry

Binding affinities of Fabs H1 and 2D2 against CHIP were determined by biolayer interferometry (Octet RED384 system). All samples were prepared in Black 384 well microplates. Biotinylated CHIP was immobilized to streptavidin tips (Sartorius), and association and dissociation steps were both performed in 1% BSA - 25 mM HEPES, 50 mM KCl. Raw data was exported using Octet BLI Analysis 12.2.2.4 and K_D_ was determined using steady state kinetics analysis on GraphPad Prism 9.

### Cryo-EM sample preparation, data collection and image processing

CHIP:Fab complex samples were produced by incubating CHIP (2 µM) and excess H1 or 2D2 (8 µM) in binding buffer (40 mM HEPES, 100 mM KCl, pH 7.4) for 3 hrs at room temperature. Additionally, WT Hsp70 (2 µM) was present in the samples. 3.5 μL of CHIP:Fab complex sample was applied to a 300 mesh graphene oxide (bare GO) coated-Quantifoil 1.2/1.3 holey carbon grid that was glow discharged (PELCO easiGlow, 15 mA, 2 min) immediately before sample application^47^. Grids were blotted using Whatman Grade 595 filter paper (GE Healthcare) for 3 s with a blot force of 0 at 4°C and 100% humidity using a Vitrobot (Thermo Fisher Scientific) and plunge frozen into liquid ethane. Samples were loaded onto a Titan Krios TEM (Thermo Fisher Scientific) operated at 300 kV and equipped with a BioQuantum K3 Imaging Filter (Gatan) using a 20 eV zero-loss energy slit (Gatan). Data was acquired with SerialEM^48^ in super-resolution mode at a calibrated magnification of 59,952x, corresponding to a physical pixel size of 0.834 Å. A nominal defocus range of −0.8 to −1.8 µm was used with a total exposure time of 2 s fractionated into 0.0255-s frames for a total dose of 66 e^−^/Å^2^ at a dose rate of 15 e^−^ per pixel per s. Movies of images were subsequently corrected for drift and dose-weighted using MotionCor2^49^.

For the CHIP:H1 complex sample, a total of 38 k micrographs were collected initially and dose-weighted sums were processed and CTF estimation was performed in cryoSPARC^50^. This was followed by blob-based particle picking, 2D classification, selection of classes for template picking, another round of 2D classification, particle selection (1.1 million particles), ab initio reconstruction, and heterogenous refinement into two identified classes of interest and a junk class. The first two classes corresponded to CHIP:H1 complexes in a 2:1 and 2:2 stoichiometric ratio. For Asymmetric CHIP:H1 2:1 complex, 482 k particles were refined using non-uniform refinement (map for ^Asym^CHIP^2^:H1^1^). Further, using a mask on H1 variable and CHIP U-box domains, focused refinement was performed to obtain a high-resolution CHIP U-box:H1 epitope focused map. 514 k particles corresponding to 2:2 CHIP:H1 complex were analyzed using 3D variability analysis (3 components) and output displayed in cluster mode (clusters=6). The 6 cluster volumes and particles were input into heterogeneous refinement for re-classification. For Asymmetric CHIP:H1 2:2 complex, 202 k particles from Classes 1 and 2 were combined and non-uniform refined (map for ^Asym^CHIP^2^:H1^2^). For Symmetric CHIP:H1 2:2 complex, 278 k particles from Classes 0, 3 and 5 were combined and non-uniform refined (map for ^Sym^CHIP^2^:H1^2^).

For the CHIP:2D2 complex sample, a total of 32 k micrographs were collected initially and dose-weighted sums were processed and CTF estimation was performed in cryoSPARC^50^. This was followed by blob-based particle picking, 2D classification, selection of classes for template picking, another round of 2D classification and particle selection (962 k particles). 9% particles showing density for full-length CHIP were refined by ab initio reconstruction and non-uniform refinement (map for full length CHIP:2D2 complex). Ab initio reconstruction of all the 962 k particles into 3 classes identified CHIP:2D2 complex (with lower TPR and U-box domain densities) in the predominant class (488 k particles) and two junk classes. Class 0 particles were further classified by heterogenous refinement into 3 classes (input Class 0 particles and Class 0,1,2 volumes). 337 k particles from first class were focused refined using non-uniform refinement with a mask around 2D2 and CHIP CC domain dimer to obtain the high-resolution CHIP CC:2D2 focused epitope map.

### Molecular modeling

To generate the model for Class 1: ^Asym^CHIP^2^:H1^1^, a homology model for human CHIP was obtained using a previous crystal structure of murine CHIP homodimer (PDB 2C2L)^13^. This, along with a Fab H1 homology model computed using SWISS-MODEL^51^ was simultaneously docked into the cryo-EM density using UCSF ChimeraX^52^ and ISOLDE^53^, followed by refinement using Rosetta Fast Torsion Relax^54,55^. Same was followed to generate the Class 2: ^Asym^CHIP^2^:H1^2^ and Class 3: ^Sym^CHIP^2^:H1^2^ models, each using two copies of the Fab H1 homology model. To generate the model for the CHIP U-box-H1 interaction interface focused map, ISOLDE^53^ and Rosetta Fast Torsion Relax^54,55^ were followed by PHENIX Real Space Refine^56^ for further modeling and refinement, and final model validation. The model for full length CHIP:2D2 complex was generated by rigid-body docking human CHIP dimer homology model and Fab 2D2 homology model computed using SWISS-MODEL^51^, using USCF ChimeraX^52^. To generate the model for the CHIP CC dimer-2D2 interaction interface focused map, ISOLDE^53^ and Rosetta Fast Torsion Relax^54,55^ were used, followed by PHENIX Real Space Refine^56^ for further modeling and refinement, and final model validation. All model statistics and density fit information are presented in Table 1 and Extended Data Figs. 3, 4, 7. The AlphaFold 3^34^ predicted models for complexes between CHIP, E2s and Ub were obtained using AF3 server (alphafoldserver.com) with inputs of 2 copies of human CHIP (residues 25-303), 2 copies of E2s (Ubc13, UbcH5A or Ube2w), and 2 copies of Ub. Models were visualized using UCSF ChimeraX^52^.

### In vitro ubiquitination assay

For the CHIP autoubiquitination assay 4X stock solutions were prepared : (1) Ube1 (800 nM) and UbcH5b (8 μM) (R&D Biosystems), (2) Ubiquitin (2 mM) (R&D Biosystems), (3) CHIP (8 μM) + increasing amounts of Fab (H1: 1x, 2x, 3x and 4x, or 2D2: 0.5x, 1x, 2x and 3x) and (4) ATP (10 mM) and MgCl_2_ (10 mM) (Sigma-Aldrich) in ubiquitination assay buffer (50 mM Tris, 50 mM KCl, pH 8.0). Ubiquitination reactions were assembled by sequentially adding 10 μL of each 4X stock solution to the reaction mixture, with ATP/ MgCl_2_ added last. The final reaction volume was 40 μL and included 200 nM Ube1, 2 μM UbcH5b, 500 μM ubiquitin, 2 μM CHIP with increasing amounts of Fab (H1: 2,4,6 or 8 µM and 2D2: 1,2,4, or 6 µM) and 2.5 mM ATP, 2.5 mM MgCl_2_. The reactions were incubated for 15 mins at room temperature, and quenched with SDS–PAGE loading buffer, heated to 95 °C, separated by SDS-PAGE, and analyzed by Western blotting using anti-CHIP (Abcam, 1:2000) antibody.

### Fluorescence Polarization Binding Assays

CHIP (± Fab) – Hsp70 C-term peptide binding assays were performed using fluorescent peptides corresponding to the Hsp70 C-terminus: 5FAM-Ahx-SSGPTIEEVD and tighter binding optimized peptide “CHIPopt”^22^ FITC-Ahx-LWWPD that were custom synthesized (GenScript). Prior to use, tracer solutions were diluted in assay buffer (25 mM HEPES, 50 mM NaCl, and 0.001% Triton X-100, pH 7.4) to a working concentration of 10 nM. Each well received CHIP ± Fab (9 µl) from a 3-fold dilution series prepared using the assay buffer. Subsequently, tracer (9 µl, 10 nM) was added to each well, resulting in a final concentration of 5 nM and a total assay volume of 18 ml. The plate was shielded from light and allowed to incubate at room temperature for 15 min. All experiments were conducted in 384-well, black, low volume, round-bottom plates (Corning; catalog number 4511). FP values, measured in millipolarization units, were recorded with a Molecular Devices Spectramax M5 plate reader at an excitation wavelength of 485 nm and an emission wavelength of 530 nm. Data analysis was carried out using agonist vs response (three parameter) model in GraphPad Prism 9.

### In vitro Tau Aggregation Assay

Tau aggregation was assessed using Thioflavin T (ThT) fluorescence assays in 384-well plate (Corning), following previously described method^12^. Briefly, tau or p-tau samples (5 μM) were incubated with CHIP ± Fab (5 µM) in PBS (pH 7.4) containing MgCl_2_ (2 mM) for 30 minutes. Control studies were performed with tau or p-tau samples (5 μM) incubated with only Fabs H1 or 2D2 (5 µM). ThT (final concentration 5 μM; Sigma) was added, and aggregation was initiated by the addition of a freshly prepared heparin sodium salt solution (MW 8,000–25,000 Da; Santa Cruz) at a final concentration of 100 μg/mL. The reactions were conducted at 37 °C with continuous shaking, and fluorescence measurements were recorded using Spectramax M5 microplate reader (Molecular Devices) with excitation at 444 nm, emission at 485 nm, and a cutoff of 480 nm. Readings were taken every 5 minutes for 24 or 48 hours. Baseline fluorescence values from ThT in tau buffer alone were subtracted to correct for background signals. Aggregation curves, corrected for baseline, were fit to the Gompertz function to determine lag times.

## Data availability

Accession codes for atomic coordinates deposited to the Protein Data Bank and Cryo-EM maps deposited to the Electron Microscopy Data Bank will be shared in peer-reviewed publication.

## Additional information

This article contains Extended Data information.

## Author contributions

A.U., D.C., E.J.C., S-W.C, S.D., A.C.T., E.T., C.M.N, J.E.G., C.S.C., and D.S., methodology; A.U., D.S., writing– original draft; D.S., C.S.C., J.E.G., supervision; A.U., D.S., C.S.C., J.E.G., funding acquisition; D.S., C.S.C., J.E.G., resources; A.U., J.E.G., C.S.C., and D.S., writing– review and editing.

## Acknowledgements

We thank Dr. F. Wang and Dr. D. Agard for providing the bare GO cryo-EM grids used in this study. This work was supported by grants from the National Institutes of Health RF1AG068125, R01NS059690, and U54AI170792 and by an Alzheimer’s Association Research Fellowship grant (AARF-22-974237) to A.U.

## Extended Data

**Extended Data Fig 1.**
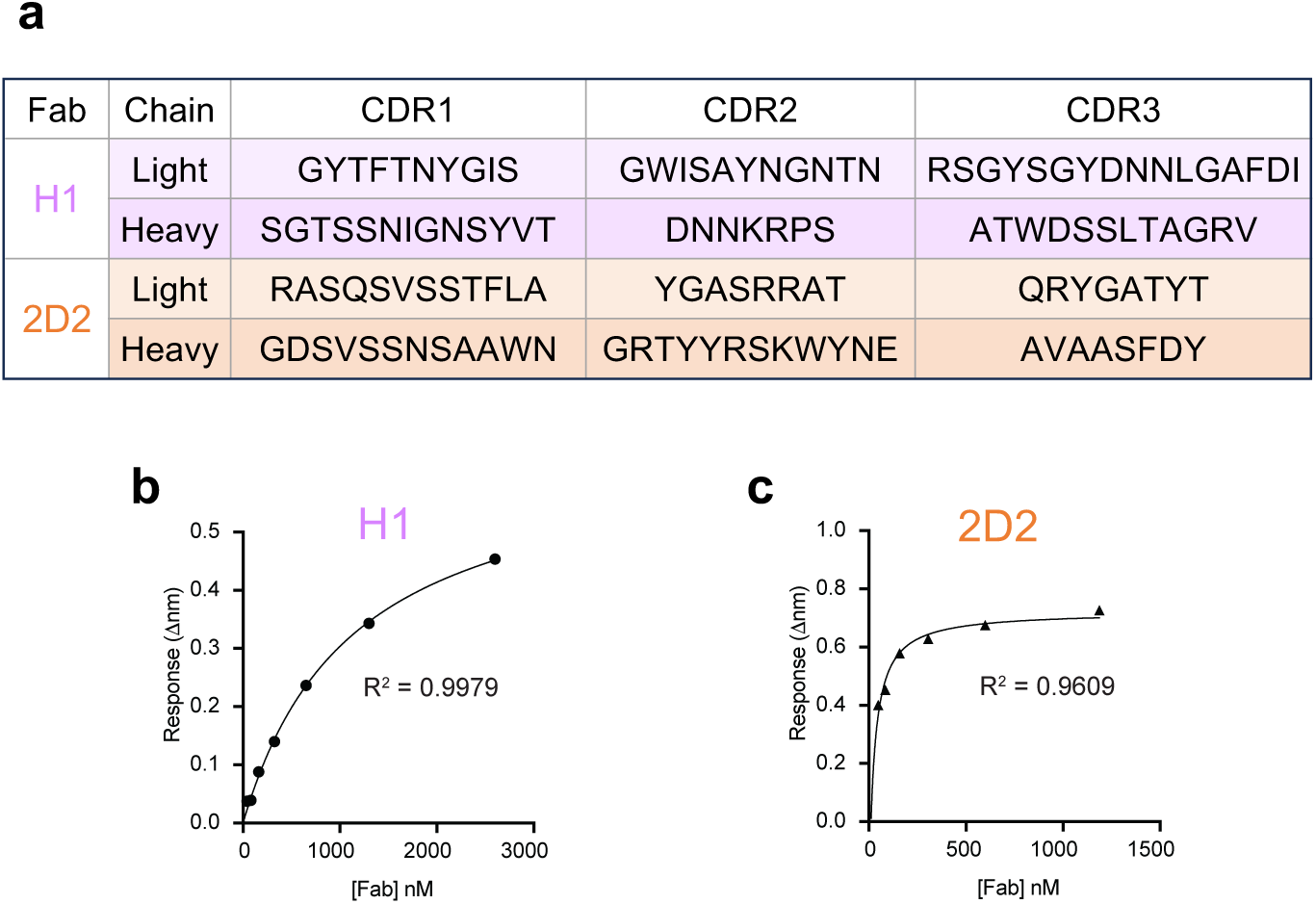
Steady state kinetics analysis for CHIP binding to H1 and 2D2. (a) CDR sequences for the light and heavy chains of Fabs H1 and 2D2 generated by biopanning. (b) Steady-state kinetic analysis of CHIP binding to Fabs was performed using BLI. Binding responses measured at varying concentrations of Fab H1 to determine the equilibrium dissociation constant K_D_ for each interaction are shown. The R² values of the curve fits are indicated to demonstrate the quality of the fits. (c) Same as (b) for CHIP binding to Fab 2D2.

**Extended Data Fig 2.**
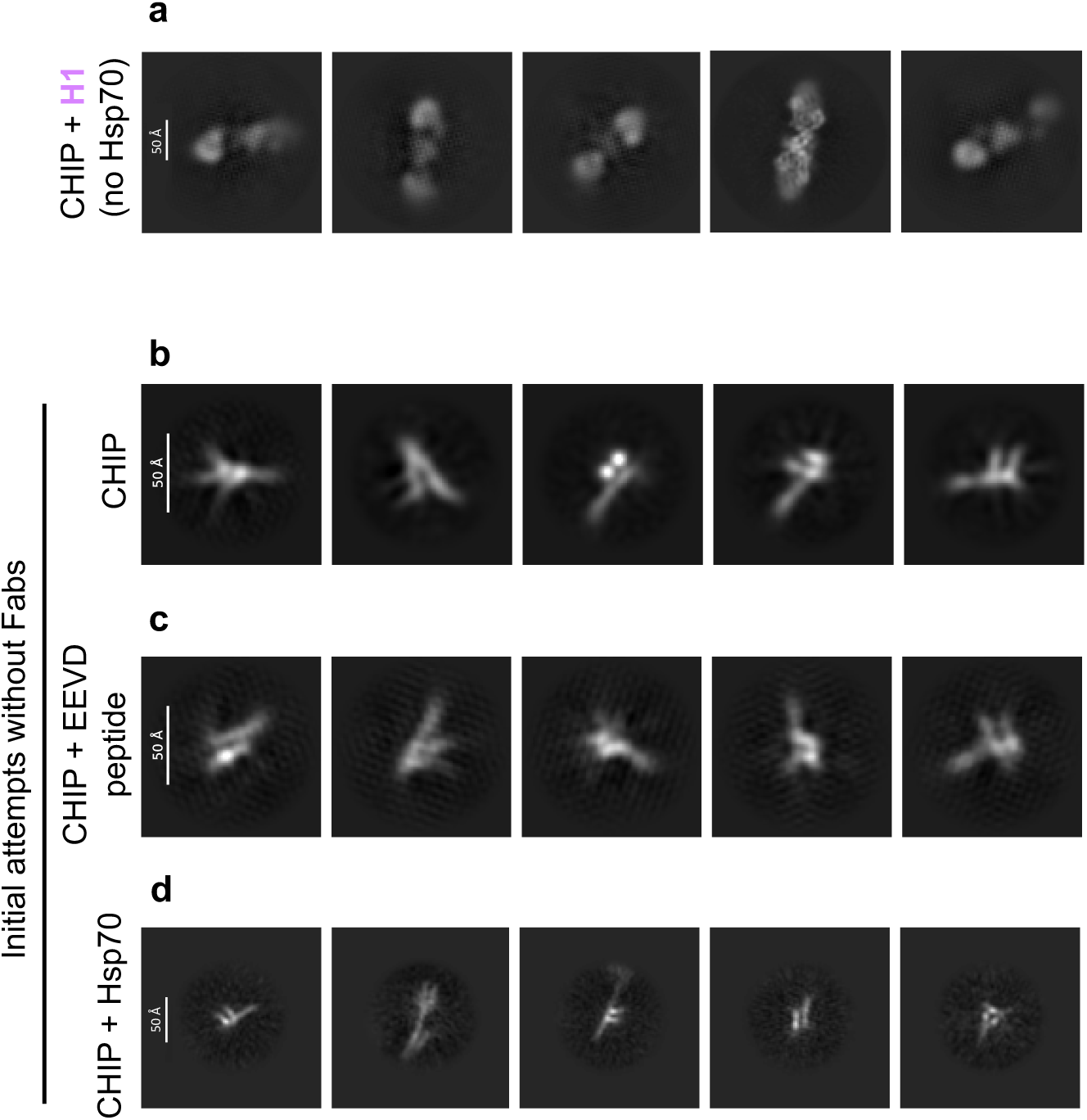
CHIP could not be resolved in initial attempts, as seen from poor 2D class averages. Representative cryo-EM 2D class averages of (a) CHIP incubated with Fab H1 showed two Fabs bound to CHIP U-box dimers, still with significant preferred orientation bias (b) CHIP alone (c) CHIP incubated with Hsp70 C-terminal EEVD peptide (SSGPTIEEVD) (d) CHIP incubated with Hsp70 (classes shown from particles ‘template picked’). Particles representing CHIP+Hsp70 complex were not observed.

**Extended Data Fig 3.**
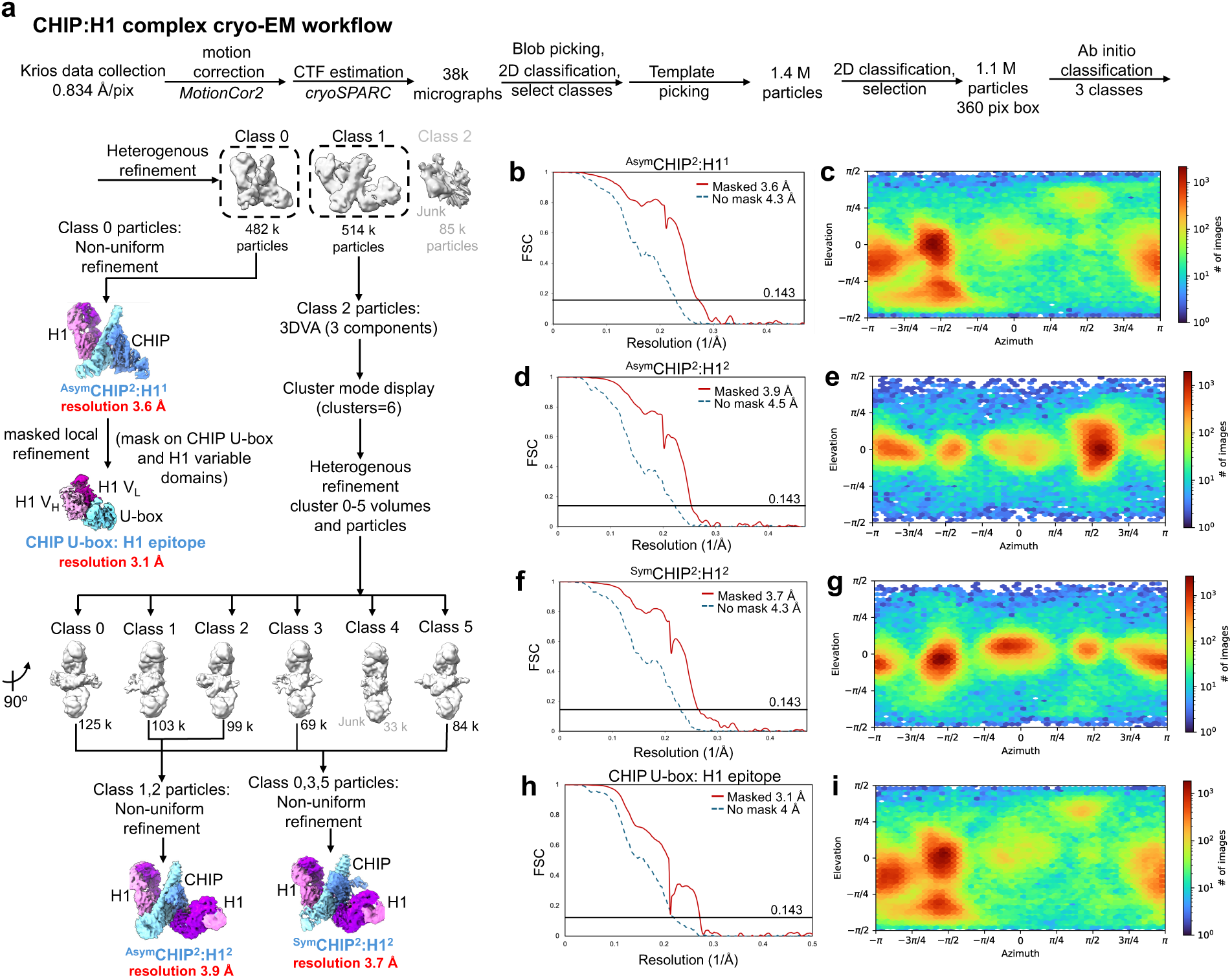
CHIP:H1 complex cryo-EM data processing, related to Figures 2,3. (a) Cryo-EM image processing workflow for CHIP:H1 2:1 and 2:2 dimer complexes. (b) Gold standard Fourier shell correlation (FSC) curves for the final masked and unmasked refinements of ^Asym^CHIP^2^:H1^1^. (c) 3D angular distribution plot of the particles used in the final reconstruction of ^Asym^CHIP^2^:H1^1^. (d) Same as (c) for ^Asym^CHIP^2^:H1^2^. (e) Same as (d) for ^Asym^CHIP^2^:H1^2^. (f) Same as (c) for ^Sym^CHIP^2^:H1^2^. (g) Same as (d) for ^Sym^CHIP^2^:H1^2^. (h) Same as (c) for refined map of CHIP U-box:H1 epitope. (i) Same as (d) for refined map of CHIP U-box:H1 epitope.

**Extended Data Fig 4.**
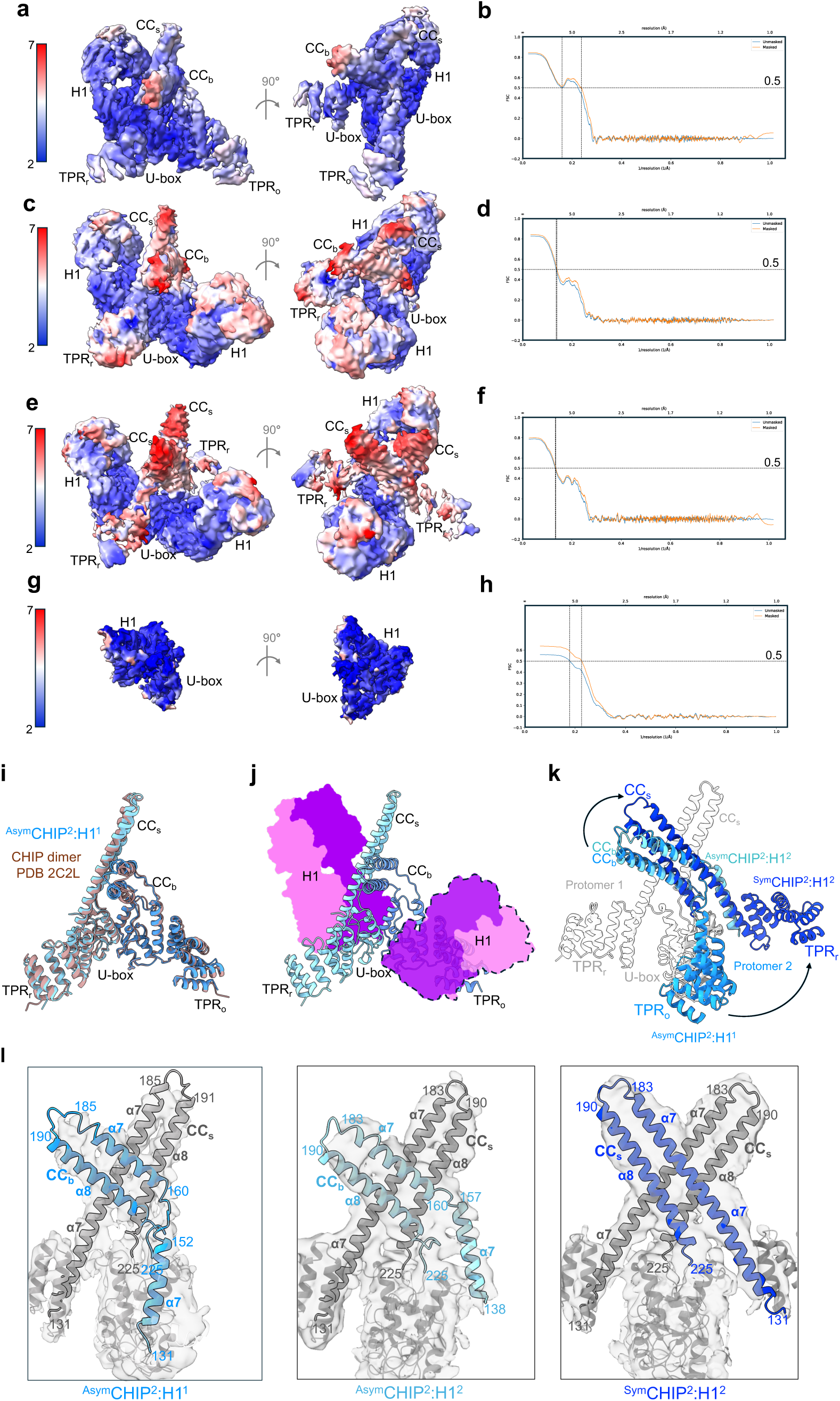
Details, overlay and comparisons of three Fab H1-stabilized CHIP dimer conformations, related to Figures 2,3. (a) Views of cryo-EM map of ^Asym^CHIP^2^:H1^1^ reconstruction colored by local resolution determined using cryoSPARC2^50^ (b) Map vs. model FSC of ^Asym^CHIP^2^:H1^1^. Phenix (Version 1.19.1) was used to determine the map-model FSC and calculate a soft mask with standard parameters^56^ (c) Same as (a) for ^Asym^CHIP^2^:H1^2^. (d) Same as (b) for ^Asym^CHIP^2^:H1^2^. (e) Same as (a) for ^Sym^CHIP^2^:H1^2^. (f) Same as (b) for ^Sym^CHIP^2^:H1^2^. (g) Same as (a) for refined map of CHIP U-box:H1 epitope. (h) Same as (b) for refined map of CHIP U-box:H1 epitope. (i) ^Asym^CHIP^2^:H1^1^ model (CHIP dimer colored blue, H1 not shown for clarity) overlaid on crystal structure of murine CHIP dimer (PDB 2C2L, colored brown). (j) Second H1 from ^Asym^CHIP^2^:H1^2^ model (shown as outlined transparent surface colored pink/magenta) overlaid on Class 1 ^Asym^CHIP^2^:H1^1^ model (CHIP dimer ribbon colored blue, single bound H1 shown as surface colored pink/magenta) showing the structural clashes between the occluding TPR_o_ domain in ^Asym^CHIP^2^:H1^1^ model and the ^Asym^CHIP^2^:H1^2^ second H1 positions, revealing displacement of occluding TPR_o_ by the second H1 in Class 2 ^Asym^CHIP^2^:H1^2^ (k) Overlay of ^Asym^CHIP^2^:H1^1^ (protomer 2 colored blue), ^Asym^CHIP^2^:H1^2^ (protomer 2 colored cyan), and ^Sym^CHIP^2^:H1^2^ (protomer 2 colored navy) models showing large domain rearrangements in CHIP protomer 2 compared with protomer 1 of ^Asym^CHIP^2^:H1^1^ (colored grey, corresponding H1s not shown for clarity) (l) Details of the differences in the CC domain arrangements (residues 131-225) in models: ^Asym^CHIP^2^:H1^1^ (protomer 2 colored blue), ^Asym^CHIP^2^:H1^2^ (protomer 2 colored cyan), and ^Sym^CHIP^2^:H1^2^ (protomer 2 colored navy), shown as models fit in low-pass filtered, zoned maps. Protomer 1 and U-box domains colored grey, occluding TPR domain of protomer 2 in ^Asym^CHIP^2^:H1^1^ shown as transparent ribbon and all H1s are not shown for clarity.

**Extended Data Fig 5.**
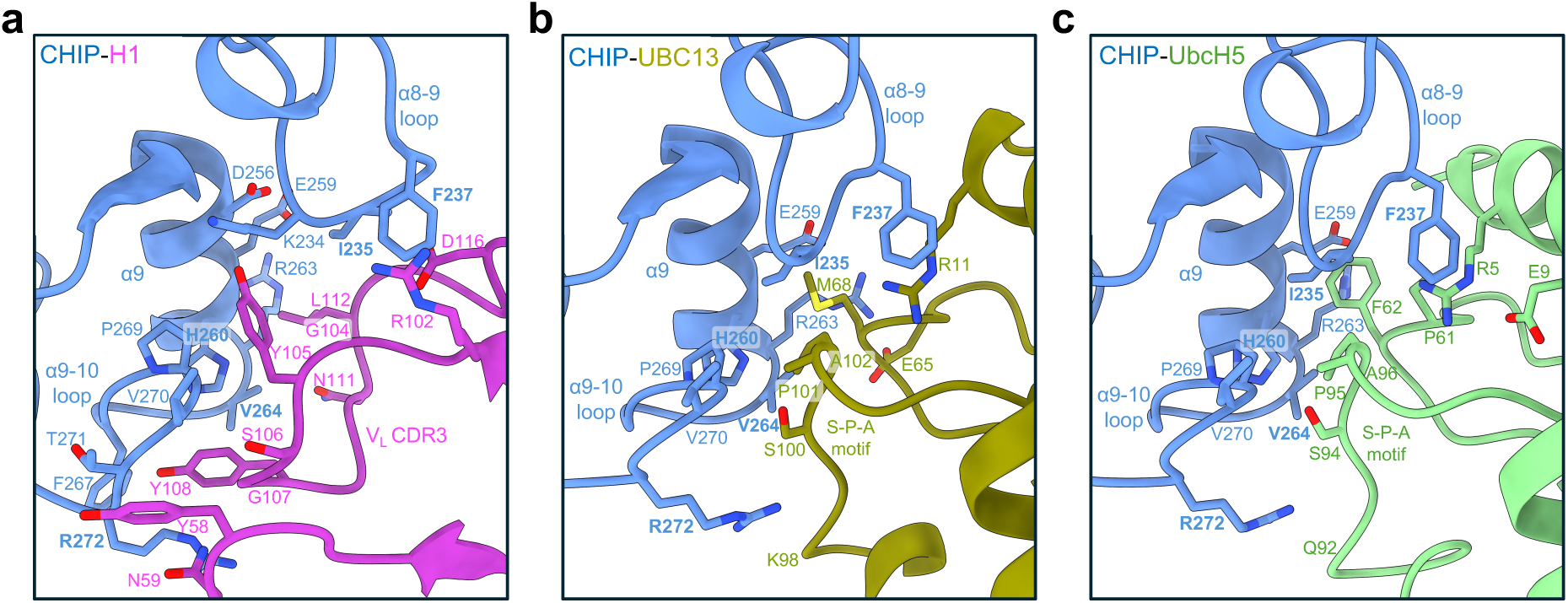
H1 binding mimics the occupancy of CHIP U-box domains by cognate E2 enzymes. (a) Details of the binding mode of H1 V_L_ with CHIP U-box residues in loops α8-9 (229-239) and α9-10 loop (264–275), and helix α9 (254–264) highlighting the hydrophobic contacts made by H1 V_L_ loops CDR3 and 2. Residues important for CHIP-E2 interactions are highlighted (bold)^13^. (b) Details of the binding mode of E2 enzyme Ubc13 with CHIP U-box residues in same loops α8-9, α9-10 and helix α9, highlighting the hydrophobic contacts made by specificity determining “S-P-A” motif residues in loop7 of Ubc13 (PDB 2C2V)^13,32^ (c) same for E2 enzyme UbcH5A interactions with CHIP U-box (PDB 2OXQ)^14^. CHIP residues in panels (b) and (c) numbered according to the human CHIP sequence.

**Extended Data Fig 6.**
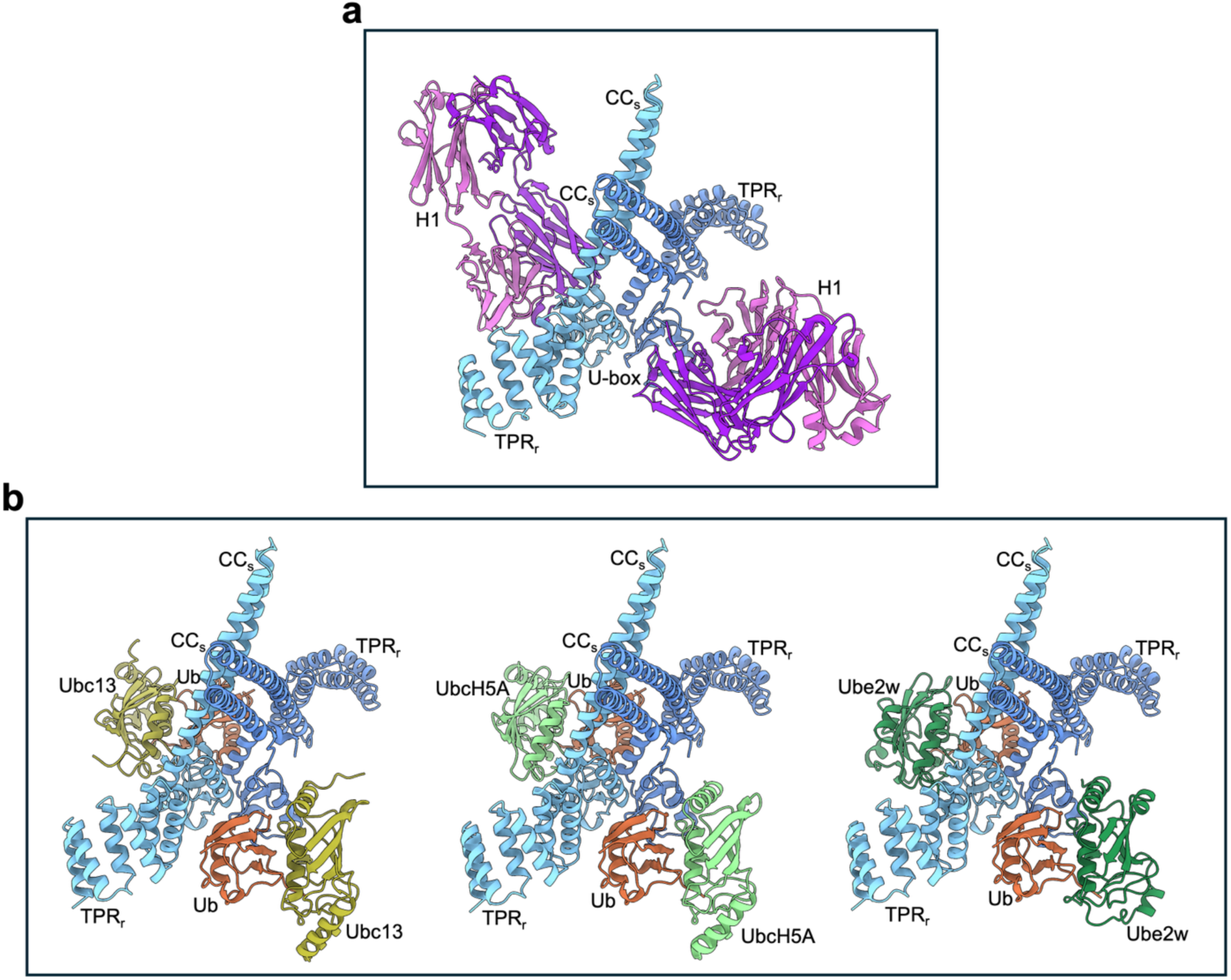
Full-site 2:2 symmetric CHIP dimer-E2-Ub binding interactions predicted by AlphaFold 3. (a) ^Sym^CHIP^2^:H1^2^ model (b) AlphaFold 3 predicted structural models for CHIP dimers interacting with E2s (Ubc13 or UbcH5A or Ube2w) and Ub, all showing symmetrical CHIP dimer conformations bound to 2 E2-Ub, predicting 2:2 full-site binding. The E2-bound ubiquitins make additional *trans-* contacts with the U-box and TPR domains of the opposing CHIP protomers. AlphaFold 3 models prediction metrics: predicted template modeling (pTM), interface pTM (ipTM) scores and global pLDDT (average of all scores): for complex with E2 Ubc13: pTM = 0.74, ipTM = 0.77, pLDDT: 85; for complex with E2 UbcH5A: pTM = 0.75, ipTM = 0.77, pLDDT: 85.60; and for complex with E2 Ube2w: pTM = 0.74, ipTM = 0.76, pLDDT: 83.85.

**Extended Data Fig 7.**
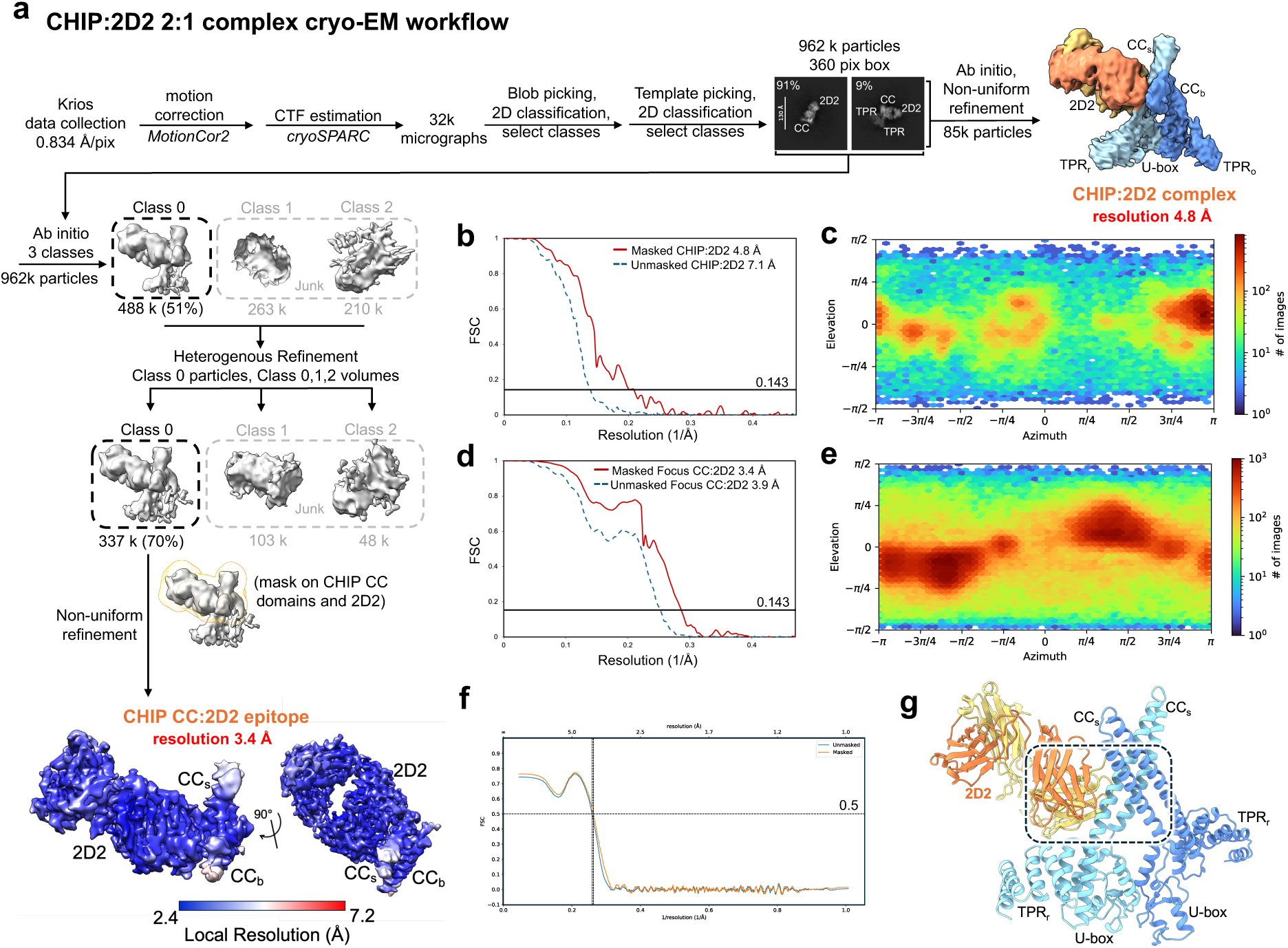
CHIP:2D2 complex cryo-EM data processing, related to Figure 4. (a) Cryo-EM image processing workflow for the CHIP:2D2 2:1 complex. Final refined map of CHIP CC:2D2 epitope is colored by local resolution, determined using cryoSPARC2^50^ (b) Gold standard FSC curves for the final masked and unmasked refinements of CHIP:2D2 complex. (c) 3D angular distribution plot of the particles used in the final reconstruction of CHIP:2D2 complex. (d) Same as (b) for refined map of CHIP CC:2D2 epitope (e) Same as (c) for refined map of CHIP CC:2D2 epitope (f) Map vs. Model FSC of refined map of CHIP CC:2D2 epitope. Phenix (Version 1.19.1) was used to determine the map-model FSC and calculate a soft mask with standard parameters^56^. (g) Overlay of CHIP CC:2D2 epitope model (2D2 colored yellow/orange, corresponding CHIP CC domains not shown for clarity) on the ^Sym^CHIP^2^:H1^2^ CHIP dimer model (CHIP dimer colored blue, corresponding H1s not shown for clarity) highlights that symmetrical conformation of CHIP dimers is incompatible for 2D2 binding to both the CHIP CC_s_ domains.

**Extended Data Fig 8.**
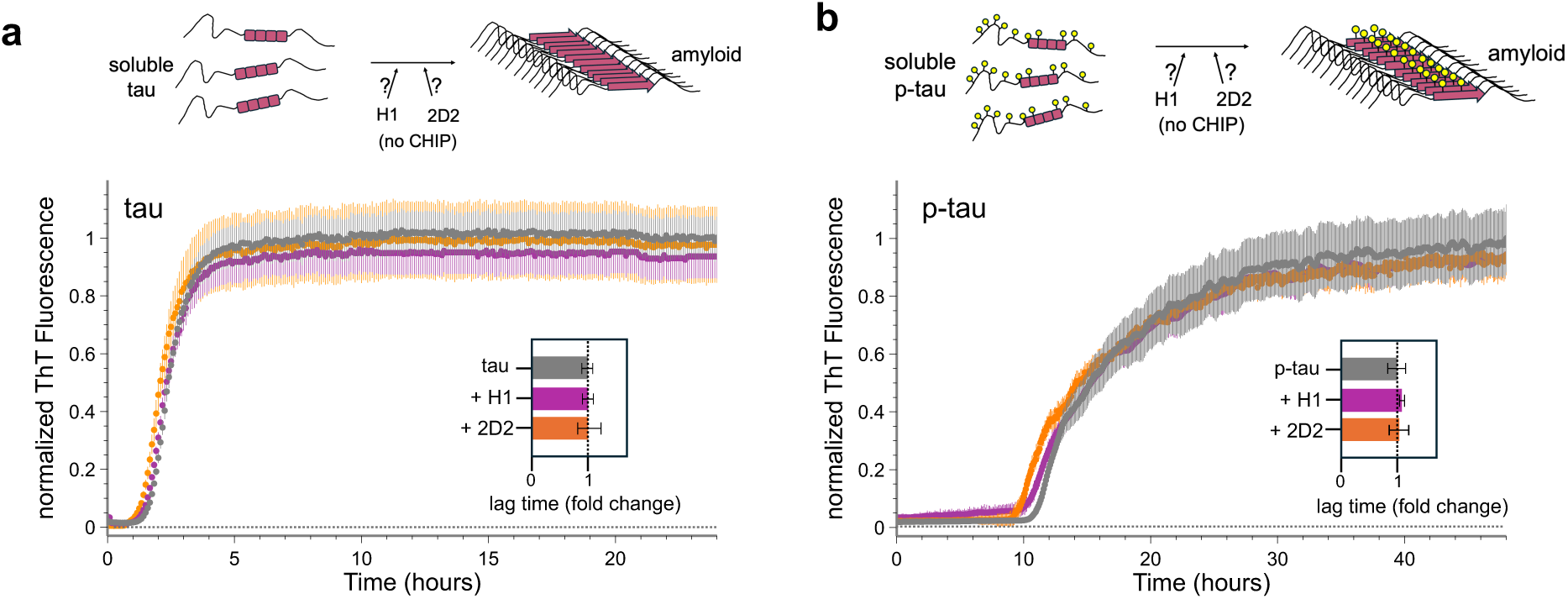
Testing effects of presence of Fabs H1 and 2D2 (no CHIP) on tau and p-tau aggregation. (a) Normalized ThT curves of unmodified 0N4R tau aggregation in the presence of Fabs H1 or 2D2. Data is shown as mean ± SD (n=3). [tau] and [Fab]= 5 μM. Derived lag time changes for unmodified tau aggregation in the presence of Fab is shown. Data is normalized to free tau and represented as mean ± SD. (b) Normalized ThT curves of 0N4R p-tau aggregation in the presence of Fabs H1 or 2D2. Data is shown as mean ± SD (n=3). [p-tau] and [Fab]= 5 μM. Derived lag time changes for p-tau aggregation in the presence of Fabs is shown. Data is shown relative to free p-tau and represented as mean ± SD.

